# Simple, reference-independent analyses help optimize hybrid assembly of microbial community metagenomes

**DOI:** 10.1101/2023.09.12.557416

**Authors:** Garrett J. Smith, Theo van Alen, Maartje van Kessel, Sebastian Lücker

**Affiliations:** Department of Microbiology, Radboud Institute for Biological and Environmental Sciences, Radboud University, Nijmegen, The Netherlands; Department of Microbiology, The Ohio State University, Ohio, USA

## Abstract

Hybrid metagenomic assembly, leveraging both long- and short-read sequencing technologies, of microbial communities is becoming an increasingly accessible approach, yet its widespread application faces several challenges. High-quality references may not be available for assembly accuracy comparisons common for benchmarking, and certain aspects of hybrid assembly may require dataset-dependent, empirically-guided optimization rather than application of a uniform approach. In this study, several simple, reference-free characteristics – gene lengths and read recruitment – were analyzed as reliable proxies of assembly quality to guide hybrid assembly optimization. These characteristics were further explored in relation to reference-dependent genome- and gene-centric analyses that are common for microbial community metagenomic studies. Here, two laboratory-scale bioreactors were sequenced with short and long read platforms, and assembled with commonly used software packages. Following long read assembly, long read correction and short read polishing were iterated to resolve errors. Each iteration in this process was shown so have a substantial effect on gene- and genome-centric community composition. Simple, reference-free assembly characteristics, specifically changes in gene fragmentation and short read recruitment, explored throughout this process replicated patterns of more advanced analyses seen in published comparative studies, and therefore are suitable proxies for hybrid metagenome assembly accuracy to save computational resources. Hybrid metagenomic sequencing approaches will likely remain relevant due to the low costs of short read sequencing, therefore it is imperative that users are equipped to estimate assembly accuracy prior to downstream gene- and genome-centric analyses.

## Background/Introduction

Though the increasing variety of high-throughput short- and long-read (meta)genomic sequencing technologies are only within their first decades of existence, both the sequencing technologies and software development have flourished and has already been implemented to study microbial ecosystems^1–15^. Integrating both short- and long-read platforms for single microorganisms or microbial communities is gaining popularity because they compensate for the other’s weaknesses – shorter reads achieve higher accuracy while longer reads offer better contiguity^7,11,16–20^. Despite the technical costs decreasing and technologies improving, hybrid strategies currently are more cost-effective for recovering high-quality Metagenome-Assembled Genomes (MAGs) from microbial communities than either platform alone^12^. However, many challenges, some biological and some technical, remain that may deter users from seeking relatively complicated approaches like hybrid (meta)genomic sequencing of microorganisms in complex ecosystems.

Hybrid (meta)genomic assembly implements multiple and/or iterative processes to help overcome the limitations of the sequencing technologies. One main strategy is to assemble short reads and then bridge gaps with long reads, leading to assemblies that are accurate but less contiguous^1,5,8,11,14,18,19^. The other main strategy is to assemble long reads and then iteratively use short reads to resolve the errors, leading to a more contiguous assembly that retains errors^1,5,8,11,14,17–26^. In the recent past and near future, as long read accuracy increases tremendously, this second strategy may become overall more favorable for the field because, intuitively, better contiguity enables better gene- and genome-resolved analyses. Most benchmarks for the second strategy have provided a general set of guidelines for implementing the iterative processes to improve upon long read assemblies: using several tools is preferred, long read correction prior to short read polishing is advantageous, and iterative processes have diminishing returns^4,12,13,17,22,26–31^. These concepts are incorporated into pipelines for automating the optimization of hybrid microbial (meta)genomic assemblies^19,29^, suggesting they are core aspects of hybrid metagenomic assembly even for microorganisms.

Intriguingly, despite the existence in literature essentially since the development of long read sequencing, the iterative processes for fixing errors in long read assemblies has been less thoroughly investigated. In particular, one challenge in performing hybrid assembly for a microbial community metagenome is identifying the number of iterations to maximize the quality of the data. Generally, there is little consistency of tool choice and the number of iterations among published studies of microbial communities or other datasets reconstructed using LR assembly^1–3,5–11,13,14,26,30^, and arguably even less consistency among benchmarking studies^3,11,14,17,18,20–24,26,27,31–43^. While most studies have determined the approach *a priori*, some evidence suggests that an unsupervised but empirically-guided approach combining various tools optimizes the hybrid microbial assemblies because certain tools are better able to fix certain errors, and some may even degrade the quality by re-introducing errors^4,17,21,26,31^. This suggests that is very likely that ideal hybrid assemblies for microbial communities may not be achieved using a universal or standard methodology, but rather likely varies depending not only on the biology of the system but also on tool choices.

An additional challenge of applying hybrid assembly to a complex microbial community is determining the quality of the assembly itself. Typically during benchmarking, assembly qualities are assessed by comparing to high-quality references, *e.g.*, count of differences (mis-assemblies), genomic alignments, or specific marker gene presences^5,12,18,22,23,30,32,33,37,44^, which can lead to poor interpretation of multiple or divergent genomes, as well as high computational costs. In the absence of high-quality reference genomes, the characteristics of an ideal hybrid assembly of a complex microbial communities are less clear. Common, reference-independent statistics for comparing assemblies, for example contig counts, total base pairs assembled, and L/N50 and similar metrics, might not significantly change at a metagenome assembly scale during iterative processes that fix relatively small-scale errors^17,26,28,30,31^. Therefore, typical assembly quality assessments are not suitable for many complex microbial community metagenomic datasets.

Benchmarking of individual bacterial genomes suggested that read recruitment, and to a lesser degree gene counts and lengths, are useful reference-free proxy indicators of assembly quality^17^. While there are advanced statistics, both of gene length distributions and read recruitment profiles are relatively simple to calculate, and often precede downstream analyses. In several cases, gene fragmentation and read recruitment profiles, often in comparison to references, have been used to asses recovered MAG quality or the accuracy of assemblies themselves^4,8,11–13,15,17,18,22,45^, but in many cases there was no apparent evaluation or optimization of hybrid assembly of microbial community metagenomes^1,2,5,9–11,14,45^. Given the challenges of applying hybrid assembly approaches for microbial community metagenomes, and the likelihood that no universal approach works best for all datasets, characteristics that are reference-independent and relative to the dataset may be the most suitable to estimate quality and empirically guide optimization.

Here, we examined multiple reference-independent and -dependent assembly characteristics in order to determine which would be effective for empirically optimizing hybrid metagenomic assembly of a uncharacterized, complex microbial communities. Two long-term laboratory-scale nitrifying bioreactors were sequenced using both Illumina MiSeq (short-read, SR) and Oxford Nanopore Technology (long-read, LR) platforms and assembled with multiple programs to allow for initial biological and computational variation. LR assemblies were then corrected with LRs and polished with SRs using prevalent tools in the field with low computation demands. Rather than assess the efficacy of tool, we sought to (1) observe the impact of iterative processes on community reconstruction, and (2) identify reference-independent assembly characteristics to help estimate assembly accuracy to optimize the approach. We first established that there is significant variation in community composition during these iterative processes as a result of the errors being fixed, but these patterns largely match published studies. We then demonstrated that simple, reference-independent assembly characteristics, for example coding gene counts and bps or SR recruitment, not only follow these same patterns but also were robustly correlated with more advanced reference-agnostic metrics, thus serving as practical proxies for assembly accuracy and community reconstruction. Finally, we offer practical guidelines to aid users for empirically optimizing hybrid assembly of microbial community metagenomes.

## Methods

The tools, their purpose, and the rationale for or advantages of their use in this study are described in Text S1, and summarized in Table S1.

### Long and short read sequencing

Biomass was collected in November 2020 and March 2021 from long-term, autotrophic nitrifying enrichment cultures maintained in either oxygen-(OLR) or nitrogen-limited (NLR) conditions in a tandem laboratory-scale bioreactor system inoculated in 2015 with wastewater treatment plant activated sludge from the Bavaria Brewery in Lieshout, The Netherlands (51.518666 N, 5.613009 E). The cultivation of these bioreactors and the reconstruction of their microbial communities is an on-going project and will be published separately in a following study. Genomic DNA was extracted from the biomass using a conventional N-cetyl-N,N,N,-trimethyl ammonium bromide (CTAB) protocol in 2020, and the Powersoil DNA Isolation kit (Qiagen) in 2021 with minimal modifications to the manufacturer’s directions to reduce DNA shearing such as inverting rather than vortexing or pipetting to mix. wo DNA isolation protocols were used because it is standard in our group due to strong evidence that it introduces sufficient bias to aid differential coverage binning^46–48^.

In total, 1 ng of DNA for both the OLR and NLR reactors was used to prepare a library using the Nextera XT kit (Illumina, San Diego, California U.S.A.) according to manufacturer’s instructions. After quality and quantity check of the libraries, they were paired-end sequenced (2x 300bp) using the Illumina MiSeq sequencing machine and the MiSeq Reagent Kit v3 (San Diego, California USA) according to manufacturer’s protocols. Oxford Nanopore Technologies (ONT) sequencing was done with 840 and 1670 ng DNA for the OLR and NLR reactors, respectively, after library preparation using the Ligation Sequencing Kit 1D (SQK-LSK108) and the Native Barcoding Expansion Kit (EXP-NBD104), according to the manufacturer’s protocols (Oxford Nanopore Technologies, Oxford, UK). The libraries were loaded on a Flow Cell (R9.4.1) and sequenced on a MinION device (Oxford Nanopore Technologies, Oxford, UK), according to the manufacturer’s instructions. Guppy (version 4.0.11)^49^ was used to basecall fast5 files using the dna_r9.4.1_450bps_hac.cfg model, both provided by Oxford Nanopore Technologies.

Raw or basecalled sequencing reads for both bioreactors and technologies are available at NCBI via BioProject PRJNA1005948 as the following BioSamples: SAMN37004618, raw MiSeq reads for the OLR; SAMN37004620, raw Oxford Nanopore Technologies reads for the OLR; SAMN37004619, raw MiSeq reads for the NLR; SAMN37004621, raw Oxford Nanopore Technologies reads for the NLR.

### Long-read, short-read, and hybrid microbial community metagenomic assembly

An overview of the experimental design is shown in Fig. 1. All computational programs were employed with default settings unless explicitly stated. Generic example code is available in File S1. Sequencing yielded 3.1-4.6 Gbp per library (Table S2) following read trimming and length and quality control using BBduk (BBtools version 37.76)^50^ with a minimum phred score 18 and length of 200 bps for the MiSeq reads, and porechop (version 0.2.3_seqan2.1.1)^51^ with minimum split length of 3000 bps for the ONT reads.

**Figure 1.**
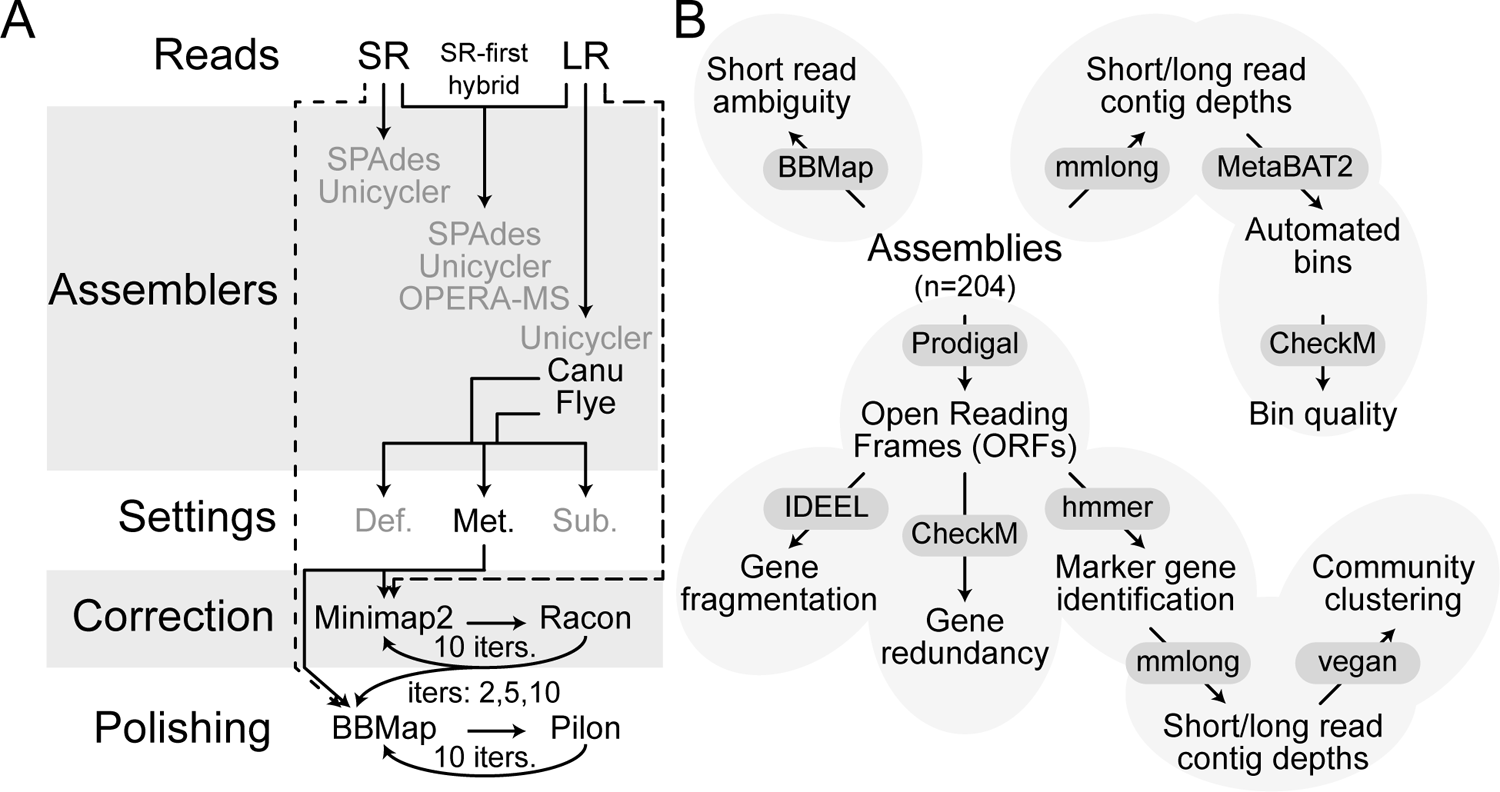
Schematic overview of assembly, correction, and polishing. (A) Assembly, correction, and polishing experimental design. Broken lines represent read recruitment steps for either correction or polishing. Acronyms: SR, short read; LR, long read; Def., default settings; Met., metagenome-optimized settings; Sub., sub-sampled LRs with metagenome-optimized settings. (B) Simplified diagram of analyses performed and the tools used to perform them for each assembly and after each iteration of correction or polishing.

Three common SPAdes-dependent programs were used to generate SR-only, LR-only, and/or SR-first hybrid, *i.e.*, short read assembly followed by connecting contigs using long read alignments, assemblies: hybridSPAdes (version 3.15.4)^40,52^ known for high-quality metagenomic assemblies, Unicycler (version 0.4.9b)^19^ with utilities for optimizing SPAdes to enable recovery of circular and high-quality single genomes, and OPERA-MS (version 0.9.0)^3^ capable of automated refinement of high-quality individual genomes within a multi-species metagenome (hybrid only). Note that Unicycler, when given only long reads, does not use SPAdes and instead uses miniasm^38^ and Racon^27^. The k-mer list generated by automated selection during Unicycler hybrid assembly was used for hybridSPAdes assemblies. While the focus here was optimizing a hybrid approach that leveraged the continuity of the long-read assemblies, these SPAdes-dependent assemblies served as a baseline for comparing SR and LR assembly of these datasets.

Two programs were used for assembly of LR data: Canu (version 1.8)^34^ that performs read correction prior to assembly and is generally thought to result in more accurate assemblies, and Flye (version 2.9-b1768)^33^ that performs correction of the assembly using the input reads. Both assemblers were used in three different ways: (1) default settings (“def”) developed for single-genomes, (2) metagenomic settings (“meta”) to improve assembly of sequences with uneven depths^2,33^, and (3) metagenomic settings for approximately even bp sub-samples (“sub”) of the reads to artificially reduce the sequencing depth and possibly uncover biological variation. Long reads were sub-sampled into 12 read pools with approximately the same quantity of bps using the “subsample” utility of Trycycler (version 0.4.1)^22^. All assemblies were filtered to a 4 kbp minimum contig length using BBtools utilities prior to any downstream analyses.

### Long read correction and short read polishing

Consensus correction of individual LR “meta” assemblies was performed using Racon (version 1.3.1)^27^ up to 10 iterations, the greatest number of iterations in a hybrid assembly approach that we identified among literature^37^. For each iteration of Racon correction, minimap2 (version 2.17-r941)^53^ was used with default settings to recruit LRs to the original assembly or previous iteration’s corrected assembly to generate the overlap information. Contigs that were not corrected were retained with the optional flag “-u”. SR pile-up polishing of LR “meta” assemblies was performed using Pilon (version 1.23)^54^ to 10 iterations after 0 (original assembly), 2, 5, and 10 rounds of LR correction. These stages were chosen to include common (0 and 2) LR correction endpoints found in literature and online guidance, and extensive (5^22^ and 10^37,55^) LR correction endpoints that would help demonstrate the maximum possible value of this process. Note that many benchmarks focus on individual microbial genomes, and particularly studies exploring the quality during iterative processes, and therefore it seemed plausible that additional iterations may aid in improving assembly quality. We did not test SR polishing after 1 iteration of LR correction because it was rarely observed in published literature or online guidelines, and essentially never when Racon was used for LR correction. For each iteration of SR polishing, BBmap (BBtools version 37.76) was used to recruit SRs to the original assembly, Racon-corrected assemblies, or previous iteration’s polished assembly, using a 97.5% identity filter and retaining all ambiguous alignments. Because we found no examples of LR correction after SR polishing and it is expected to re-introduce errors, this was not performed.

### Assembly characteristics determination

Programs for estimating quality were used with default settings except when explicitly stated otherwise. Assemblies were compared using metaQuast (version 5.0.2)^56^ without references to examine the distribution of contig counts and lengths. Recruitment of SRs to the assemblies to quantify aligned reads and ambiguity was performed using BBmap (BBtools version 37.76) with retaining only perfect mappings of paired reads and ambiguous mappings randomly assigned. Open reading frames (ORFs) were predicted using Prodigal (version 2.6.3)^57^ using the “meta” procedure for quantification of gene counts and lengths, and as input for downstream analyses of assembled genes. The “meta” procedure of Prodigal was implemented prior to downstream analytical pipelines because it led to increased marker gene recovery estimates, though following similar trends (data not shown), compared to the outputs of the analytical pipelines that assume inputs are single genomes and therefore run prodigal in “single” mode by default. Phylogenetic marker gene recovery, fragmentation, and redundancy were estimated by Benchmarking Universal Single-Copy Orthologs (BUSCO, version 5.1.2)^58^ using “genome” mode and only the “bacteria_odb10” lineage, as well as CheckM (version 1.1.3)^59^ using the taxonomy workflow for the domain “Bacteria”. Gene fragmentation of entire assemblies was estimated by comparing to a DIAMOND (version 0.9.31)^60^ database of the uniref50 (downloaded 2023-03-01)^61^ dataset using IDEEL (downloaded 2023-05-13)^13^. To more concretely associate potential complete genome contigs with the domain bacteria, circular and long (>1 Mbp) contigs were also analyzed individually using the Genome Taxonomy DataBase toolkit (GTDB-tk, version 1.6.0 with reference database version r202)^62^. Based on the GTDB-tk output for circular contigs alone, there were members of at least 6 and 5 distinct Bacterial phyla present in the OLR and NLR, respectively (data not shown). Microbiomes with abundant Archaea or Eukaryotes may need to adjust these taxonomically constrained pipelines to better suit their ecosystem.

### Automated binning and beta diversity

Programs for automated binning and estimation of genome quality, as well as identification and read depth calculation of marker genes, were used with default settings except where noted otherwise. Both LRs and SRs from both bioreactor datasets were used for read depth calculation of contigs using the mmlong (version 0.1.2)^63^ utility *readcoverage2*. Automated binning using composition (*i.e.*, tetranucleotide frequency) and coverage (*i.e.*, read depth) was then performed using Metabat2 (version 2.12.1)^64^. The quality of automated bins was estimated using CheckM (version 1.1.3)^59^, with cutoffs of >50% completeness and <10% contamination scores as “medium quality” (MQ). These oversimplified, and somewhat low, thresholds are no longer *en vogue*, but represent computationally simple, rapid, and bulk estimates compared to contemporary, thorough requirements that include other information like rRNA presence and tRNA counts^65^.

The RNA polymerase subunit B (*rpoB*) is a protein-coding gene typically found only once in a genome and is universally conserved among Bacteria, Archaea, and Eukarya, and therefore is tractable to serve as a marker gene for individual species to complement the often poorly assembling 16S rRNA gene for gene-centric phylogenetic analyses of metagenomes. *RpoB* genes were identified in assembled contigs using hmmer (version 3.1b2)^66^ with the available model and thresholds for the Protein FAMily (pfam) identifier PF04563.15 (downloaded 2020-06-09). To observe beta-diversity, *rpoB*-containing contigs throughout the iterative correction and polishing processes were compiled for the each bioreactor and assembler separately, and then the LRs and SRs from both bioreactor datasets were used for read depth estimation for *rpoB*-containing contigs using the mmlong (version 0.1.2)^63^ utility *readcoverage2*. The Bray-Curtis abundance-rank dissimilarity between each read set was calculated for subsequent two-axis Non-Metric multi-Dimensional Scaling (NMDS) performed using vegan (version 2.5-7)^67^ in R (version 4.1.2)^68^, and the values were then manually mean-centered and scaled for better comparability. These NMDS analyses were unable to converge after 50 tries due to low stress, but here the species scores were used to view the clustering and trajectories of the *rpoB*-containing contigs’ “species scores” for each assembly rather than the four reads’ “site scores”.

### Further data analysis and visualization

Logs and outputs were mined for data using various bash commands. Data were ultimately imported into R (versions 4.1.2)^68^ for analysis and visualization relying primarily on tidyverse (version 1.3.1)^69^. Most calculations were additionally normalized to contig length in Mbps to make bioreactors and assemblers more comparable. No specific code was developed for this project due to the focus on applying end-user tools.

## Results

### Short read, long read, and hybrid assembly baselines

SR sequencing yielded 7.56 million paired-end reads totaling 4.06 Gbp, and 8.73 million paired-end reads totaling 4.60 Gbp of data after low quality read removal and base trimming for the OLR and NLR, respectively. LR sequencing yielded 318 thousand reads totaling 3.13 Gbp, and 416 thousand reads totaling 4.01 Gbp of data after low quality read removal and base trimming data for the OLR and NLR, respectively (Table S2). Reads were either assembled alone (SR-alone or LR-alone) or used as inputs for automated SR-first hybrid assembly. There were substantial differences between assemblies of the bioreactors of approximately 63 total assembled Mbp on average (Fig. S1), which we infer was due to the greater sequencing depths for the NLR compared to the OLR rather than any biological differences.

In addition to biological and/or sequencing depth effects on the assemblies, the assembly approach implemented by the different programs had a substantial effect on the assembly outcome. This was apparent across many aspects of these assemblies: LR assemblies were larger in terms of total bps (on average 1.35-fold), they were less fractured such that a greater fraction of bps were on larger contigs (1.69-fold), and they yielded more circularized contigs (2.34-fold for count and 5.5-fold for total bps) (Text S1, Figs. S1, S2A, S3). However, probable complete microbial genomes, *i.e.*, a single contig greater than 0.5 Mbp and ideally circular, reconstructed with SR and SR-first hybrid assemblies tended to have higher completeness scores on average (1.73-fold for long contigs and 1.38-fold for circular, Text S1, Fig. S2B, Fig. S3). Thus, baseline assemblies recapitulated most benchmarking studies showing that LR assemblies lead to better contiguity, but SR assemblies tend to be more accurate^1,5,8,11,17,21^. Still, the contiguity of the LR assemblies offers advantages for microbial community reconstruction with sufficient LR correction and/or SR polishing to resolve mis-assemblies.

From here, we explored these metagenome-optimized LR assemblies over the iterative LR correction and SR polishing iterations to ultimately determine the ideal assembly for these datasets with empirical information. We assessed the microbial composition and its variability using standard gene- and genome-centric analyses like beta-diversity and automated bin recovery. We then further analyzed characteristics that should noticeably change, in contrast to for example the size and contiguity of the assembly, and that ideally would also be reference-free: reduction in gene fragmentation because errors in LR and their assemblies lead to frameshifts that fracture genes, and increase in SR recruitment because these more high-accuracy reads should recruit to more accurate assemblies. Additionally, we followed automated bin recovery and single phylogenetic marker gene beta diversity to see how well these simple, reference-free characteristics may serve as indicators of community reconstruction.

### Marker gene beta diversity

Complementing genome-centric approaches, many studies of microbial communities applying hybrid metagenomic assembly performed gene-centric analyses^2,5–10,13,14^, often as a means of inferring metabolic capabilities and therefore ecosystem roles, but can also help access less well assembled community members to better assess extant diversity. While genome recovery has been examined as proxies for microbial metagenomic assembly accuracy, there is substantially less effort put into examining the remainder of the community members and how this might impact downstream analyses of the communities. The RNA Polymerase subunit B (*rpoB*) gene is a phylogenetic marker for all domains of life that was used here for beta-diversity clustering analysis by using their SR recruitment profiles via Non-Metric multi-Dimensional Scaling (NMDS).

As expected given the issues with LR assembly, error-fixing substantially improved marker gene recovery but surprisingly decreased read depth. Initial LR assemblies and assemblies with LR correction alone contained fewer (2 total genes) and more fractured (1/3 the mean length) *rpoB* genes compared to assemblies after the first few iterations of SR polishing (Text S1, Fig. S4). While read depths stabilized due to SR polishing, they unexpectedly decreased on average (Text S1, Fig. S5). We speculate that this phenomenon was caused by the identification of additional *rpoB* genes on low abundance contigs once SR polishing made these genes sufficiently intact to be identified.

Both increased *rpoB* recovery and changes to read depth profiles had clear effects on the trajectories of the communities during error-fixing processes, thus influencing apparent beta-diversity. Assemblies with only LR correction were noisy and remained distant from one another, compared to the convergence observed due to the first few SR polishing iterations (Fig. 2). In most cases, assemblies without LR correction were distant from assemblies with both LR correction and SR polishing, highlighting the beneficial impact of LR correction that may only be apparent after SR polishing (Fig. 2). Unfortunately, also in most cases, communities that converged during SR polishing were still distinct from those with different preceding LR iterations, indicating that this impacts the community’s even with SR polishing (Fig. 2). These results show that iterative LR correction and SR polishing processes ultimately impact downstream community reconstruction analyses.

**Figure 2.**
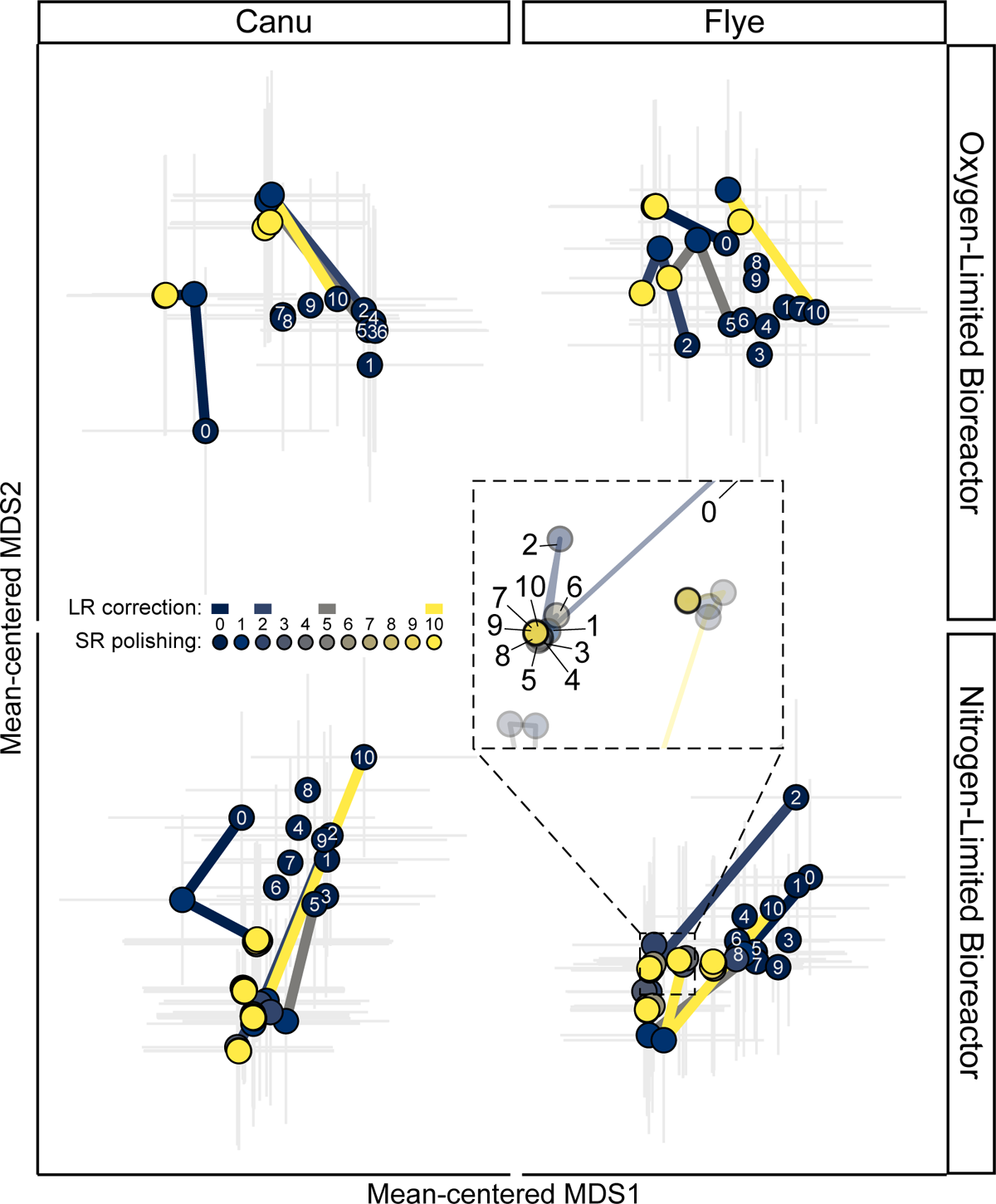
Beta diversity estimated from RNA Polymerase subunit B (*rpoB*) gene profiles for each bioreactor and Long Read (LR) assembly throughout the iterative LR correction and Short Read (SR) polishing processes. The two bioreactors are separated over vertical panels, the two LR assemblers are separated over the horizontal panels. Points are the mean-centered species scores for all rpoB genes per assembly, with gray lines showing the standard errors of the mean. Each point is colored by the SR polishing iteration, with colored lines connecting points to the preceding LR correction and subsequent SR polishing iteration. White text over points indicates the LR correction iteration prior to any SR polishing. Inset shows the overlapping points of the second LR correction stage as the communities converge due to SR polishing for the Nitrogen-Limited Bioreactor’s Flye assembly.

### Automated bin recovery

A primary goal for many microbial community metagenomic sequencing approaches is to reconstruct high-quality, ideally essentially complete genomes of as many of the members as possible^1–10,12,13^. Hybrid assembly of microbial metagenomes has already been employed to recover microbial genomes from a variety of ecosystems from waste sludge to human guts, and more^1–10,12,13^. To maximize the value of the genomes, the assembly itself needs to be accurate because both binning of contigs into genomes and their resulting quality, estimated via marker gene identification, depends on the accuracy of the assembly. In some cases, genome recovery has already been used to demonstrate or compare the accuracy of LR and/or hybrid assemblies of microbial (meta)genomes^1,7,10,15,18,33^. Here, assemblies at each stage of LR correction and SR polishing were automatically binned, and both the assemblies and bins were assessed with CheckM to estimate the completion of microbial genomes and estimate the redundancy by counting the 104 bacterial marker genes and reporting the copy number of each gene and the estimated completion and contamination percentages.

Error-fixing of LR assemblies was required for genome recovery, and in particular SR polishing greatly improved automated bin recovery. Initial assemblies led to the recovery of few medium-quality or better (MQ) automated bins, only 9.5 on average (Text S1). By following contigs across bins, we noted that binning prior to LR correction and SR polishing led to worse automated binning outcomes compared to binning after iterative error-fixing processes (data not shown). LR correction increased the number of MQ bins by three to 12.3 on average, but SR polishing alone almost doubled the number of MQ bins, and in combination with LR correction more than doubled the initial MQ bin count on average (Fig. 3, Text S1, Fig. S6). Additionally, the mean completeness scores for MQ bins remained in the low 70% range after only LR correction but increased to over 80% as a result of SR polishing (Fig. 3, Text S1, Fig. S6), indicating an increase in the quality of recovered genomic contents by combining LR correction and SR polishing. Furthermore, the substantial increases in MQ bin count and quality due to SR polishing were matched by large increases in the quantity of both number of contigs (7.17-fold) and bps (1.92-fold) contained in MQ bins (Fig. 3, Text S1, Fig. S6), indicating that SR polishing improved the ability for contigs to be binned. However, automated bin recovery metrics remained somewhat noisy and often never reached a clear plateau (Fig. 3), suggesting that genome recovery may not be a robust assembly quality indicator. In summary, automated bin analyses suggest that the iterative LR correction and SR polishing processes strongly influence genomic recovery.

**Figure 3.**
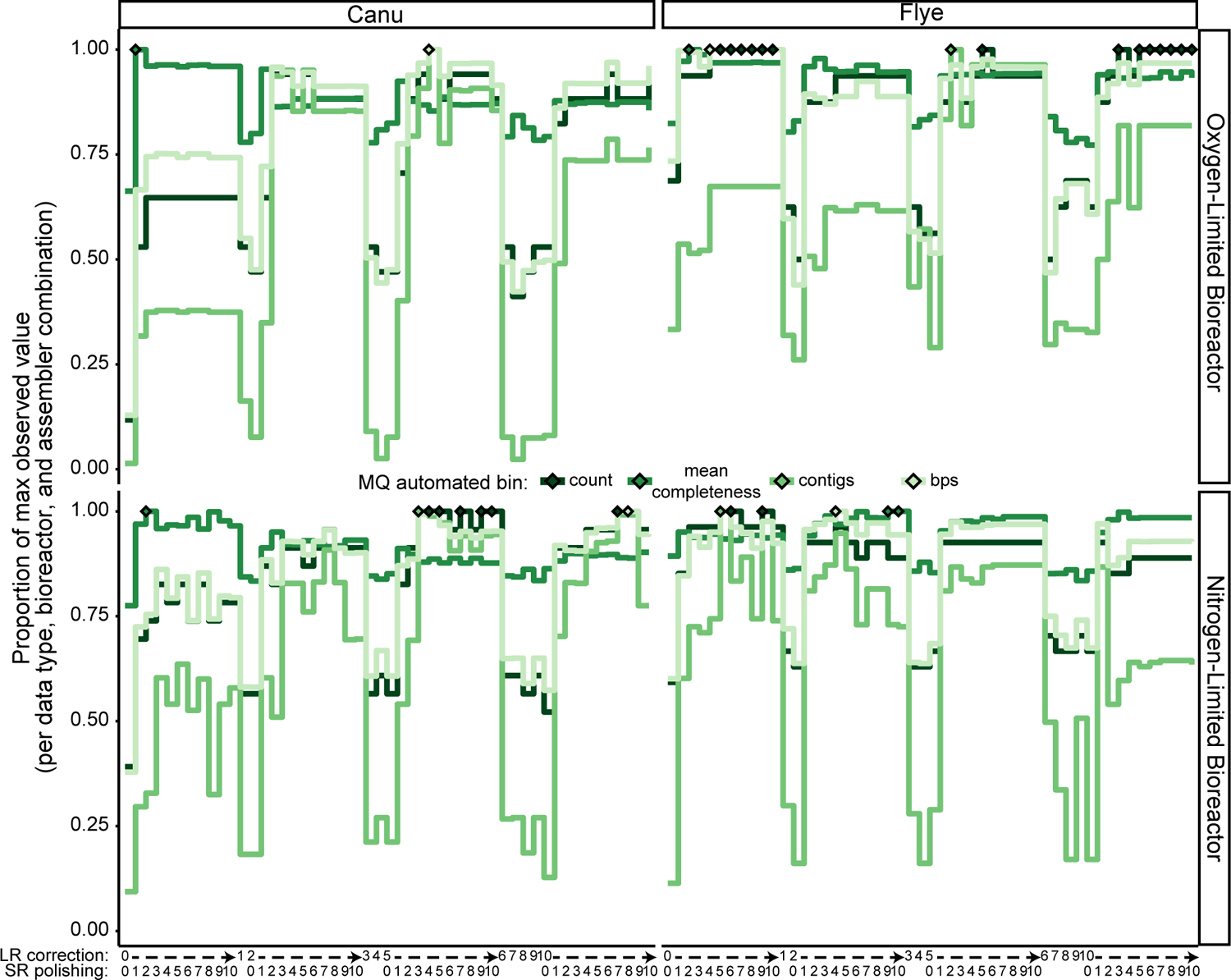
Stair plots of automated medium-quality or better (MQ) bin recovery for each bioreactor and Long Read (LR) assembly throughout the iterative LR correction and Short Read (SR) polishing processes. The two bioreactors are separated over vertical panels, the two LR assemblers are separated over the horizontal panels. The LR correction and SR polishing iterations are spread across the x-axis so that the ten SR polishing steps are immediately to the right of the respective preceding LR correction step. Each colored line represents the proportion of the maximum observed value per combination of bioreactor and assembler of one of the following characteristics MQ automated bins: the number of MQ bins recovered (“count”), the mean completeness score of the bins estimated by CheckM (“mean completeness”), the number of contigs in MQ bins (“contigs”), and the quantity of bps in MQ bins (“bps”). Matching color diamonds indicate the LR correction and/or SR polishing stage with the maximum observed value for each characteristic.

Bacterial marker gene redundancy estimated for the full assemblies followed similar patterns as automated bin recovery, and therefore may serve as a proxy to save computational resources. In contrast to bin recovery, marker gene recovery showed more clearly curves that saturated throughout the iterative error-fixing processes (Fig. S6). The estimated copy number of marker genes, as well as redundancy, *i.e.*, the sum of CheckM completion and contamination scores, reached their maximum values after two iterations of SR polishing, with greatest redundancy after both LR correction and SR polishing (Fig S6, Text S1). In fact, the redundancy of the assemblies was strongly correlated to the number of MQ bins recovered (adjusted R^2^ ≧0.85, p <<0.05, Fig. S7, Text S1). Intriguingly, slope of these regressions was approximately 0.01 (Fig. S7), or 1 genome per CheckM score of 100%, implying a near equal relationship between marker gene redundancy and complete genome recovery. However, the distance from an idealized line (Fig. S7) suggests that this hybrid assembly approach was still unable to reconstruct the genomes of all members of the microbial community. Yet, given the correlation with MQ bin recovery but an obvious plateau in redundancy, assembly-level marker gene recovery could serve as a preliminary indicator of genomic yield during the iterative processes without the need for automated binning.

### Gene fragmentation

Small scale errors (*i.e.,* insertions, deletions, and, to a lesser extent, substitutions) in LRs are known issues for using these platforms that cause gene fragmentation in the resulting assemblies^7,12,13,17,20,23,26,30,31,45,54,70^. Theoretically, as errors are fixed during the iterative LR correction or SR polishing processes then gene fragmentation should decrease, *e.g.*, fewer genes and more bps in them, and ultimately stabilize. In benchmarking studies, this is often indirectly measured by comparing differences in gene counts or marker gene inventories (*i.e.*, completion scores), or have already demonstrated that directly assessing gene lengths works as a reasonable proxy for the accuracy of hybrid assemblies of microbial (meta)genomes^4,7,11–13,17,18,22,37,45^. As a precursor to many downstream analyses, coding gene identification with programs like Prodigal allow direct estimates of coding gene lengths, while other programs like IDEEL and BUSCO can estimate gene fragmentation by comparison to reference databases and are commonly used by benchmarking or other comparative studies^4,12,13,22,30,37,71^.

Gene fragmentation substantially improved and plateaued during iterative error-fixing processes. Initial LR assemblies varied due to both bioreactor and assembler, consistent with the baseline information (Text S1). On average, gene fragmentation did not markedly improve, either in reducing the number of coding genes or increasing the quantity of bps in coding genes, due to LR correction alone (Text S1, Fig. S8). However, it did substantially improve after just two iterations of SR polishing (0.81-fold for gene count, 1.12-fold for bps in genes) after which these assembly statistics appeared saturated (Text S1, Fig. S8, Fig. S9). While small, there was a non-negligible advantage to LR correction preceding SR polishing (Text S1, Fig. S8, Fig. S9). The same patterns were noticeable not only for the entire assembly, but also for several relevant assembly fractions principally ready for downstream analyses – circular and long contigs that may represent essentially complete microbial sequences, and medium quality or better automated bins (Text S1, Figure S8, Fig. S9). As expected based on both the theory behind the iterative error fixing processes and most published studies assessing this information^4,11–13,17,18,22,37,45^, gene fragmentation was improved primarily due to the first few iterations of SR polishing.

Comparison of coding genes agnostically to reference databases showed similar patterns. After just two iterations of SR polishing, assessing the entire assemblies with BUSCO indicated that bacterial marker genes were no longer found only fragmented (Text S1, Fig. S9), and IDEEL indicated that the number of coding genes within 5% of their most similar reference sequence’s length considerably increased (2.7-fold, Text S1, Fig. S10). All gene length analyses followed similar patterns, and linear regression demonstrated that the number of coding genes within 5% of their most similar reference gene length was strongly correlated to the total number of base pairs in coding genes (R^2^ ≧0.94, p-adj <<0.05, Fig. 4). We acknowledge that the resulting data were unevenly distributed across both dependent and independent variables in this analysis because of the large differences due to, and the inflated sample sizes of, SR polishing but the correlations hold when looking at only the first three SR polishing iterations as well, albeit weaker (adjusted R^2^ 0.6-0.75, p-value <<0.05, data not shown). Additionally, similar reference-independent assembly characteristics including total number of coding genes, as well as median count of coding genes and bps in coding genes normalized to contig length, were also correlated (adjusted R^2^ >0.7, p-value <<0.05, data not shown). Therefore a less-resource demanding analysis such as gene length summarization alone is a practical proxy for more demanding analyses like gene fragmentation.

**Figure 4.**
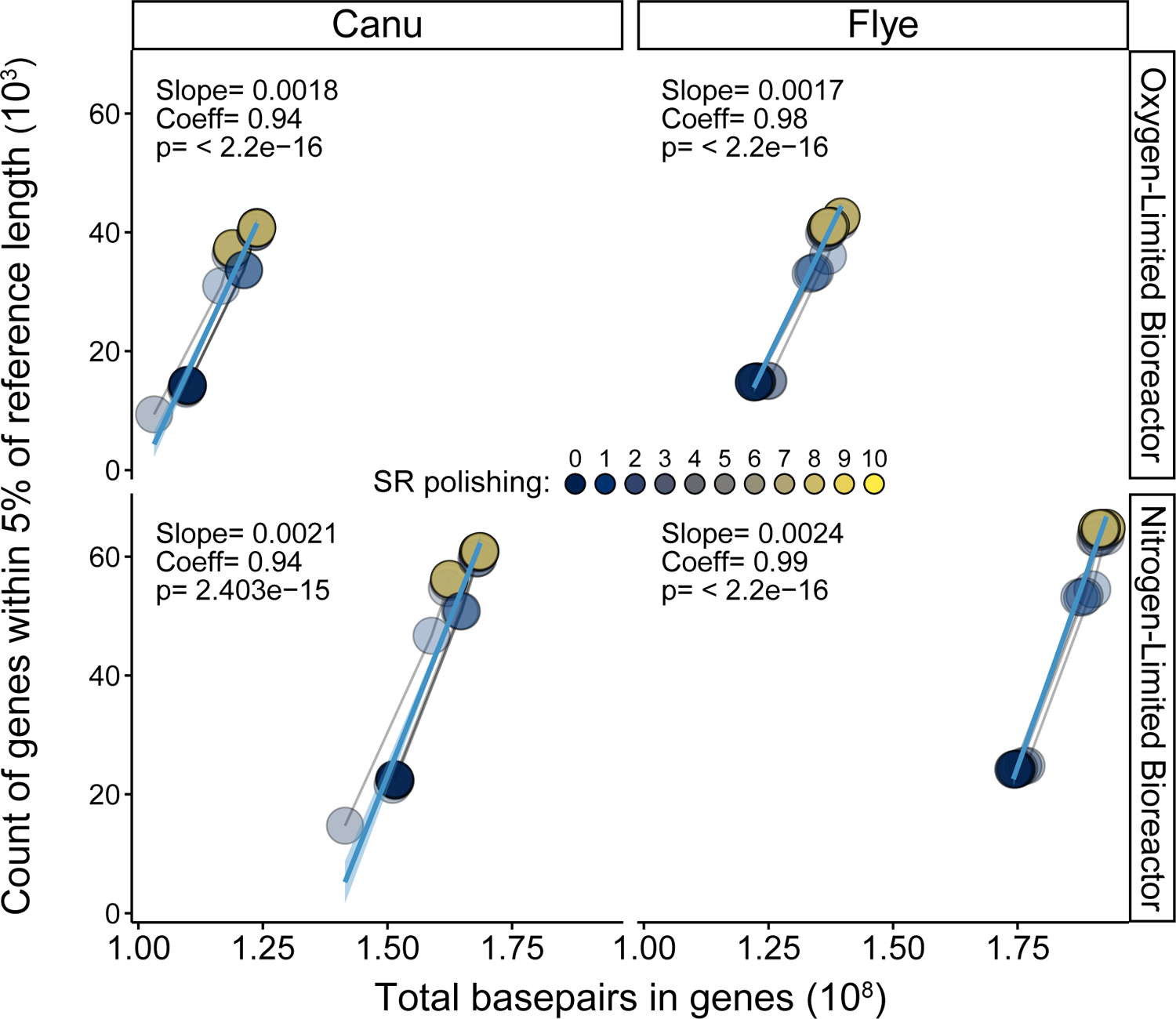
Linear correlation between calculated bps in genes and the number of these genes estimated to be similar length to reference database entries for each bioreactor and Long Read (LR) assembly throughout the iterative LR correction and Short Read (SR) polishing processes. The two bioreactors are separated over vertical panels, and the two LR assemblers are separated over the horizontal panels. Each point is colored by the SR polishing iteration, with lines connecting the points from the same LR correction iteration, all of which are partially transparent. The light blue solid lines and shaded regions are the linear regressions for the displayed data and their 95% confidence intervals. Slopes, correlation coefficients (adjusted R^2^) and p-values are displayed in upper-left corner of each panel.

### Short read recruitment

Multiple iterations of LR correction and SR polishing are often performed in benchmarks and published studies because it is expected that newly fixed errors allow different read pools to align. Then theoretically, SR recruitment would reach a maximum, stable pool as the assembly approaches high accuracy because fewer errors are fixed each iteration leading to fewer changes in read alignments. Several studies have already shown a relationship between microbial (meta)genome quality and read recruitment^4,11,12,15,17,22,45^. Additionally, in a community where multiple strains co-exist and/or species encode duplicated genes, the quantity ambiguously recruited reads (*i.e.*, aligning to two or more positions) should decrease as the minor sequence differences are resolved. Here, information-rich outputs from BBmap were analyzed for the total number of aligned SR bps, assembled contigs to which no SRs aligned, and ambiguously aligned SRs to observe the magnitudes in which these features change during LR correction and SR polishing.

Consistent with expectations and gene fragmentation results, increases to SR recruitment increased greatly due to SR polishing and then saturated after the first few iterations. The bioreactor had little apparent effect on SR recruitment to the initial assemblies, in contrast to the assembler program and (Text S1). These data, along with the larger proportion of SRs that aligned ambiguously (2.02-fold, Text S1), suggests that the Flye assemblies may be of higher quality or possibly capable of reconstructing similar yet distinct strains. On average, LR correction led to approximately 25-35% increases in total SR bps recruited, SR bps recruited normalized to contig length, and SR alignment ambiguity, but only about a 10% reduction in contigs lacking SR recruitment (Text S1). In contrast, SR polishing greatly improved SR recruitment across all measured features, including increased total SR bps recruited (3-fold) and median SR bps recruited normalized to contig length (10-fold), fewer contigs lacking any SR recruitment (30% reduction), and a smaller fraction of ambiguously aligned SRs (26% reduction, Fig. 5, Text S1). There was a clear plateau in SR recruitment by 3 iterations of SR polishing, and a noticeable improvement due to LR correction but only after some SR polishing. These observed trends match common implementation of at least 2 SR polishing iterations^2,9–13,15,17,22,29,33,45,54^. The saturation patterns of SR recruitment also appeared consistent with the decreases in changes made during SR polishing (Fig. S11), strengthening the link between using SR to improve assembly accuracy and SR recruitment profiles themselves. In summary, SR recruitment followed similar trends as all other assembly characteristics, and as a quick and common predecessor to many downstream analyses, offers a practical estimation of assembly accuracy during these iterative processes to help empirically determine their endpoints.

**Figure 5.**
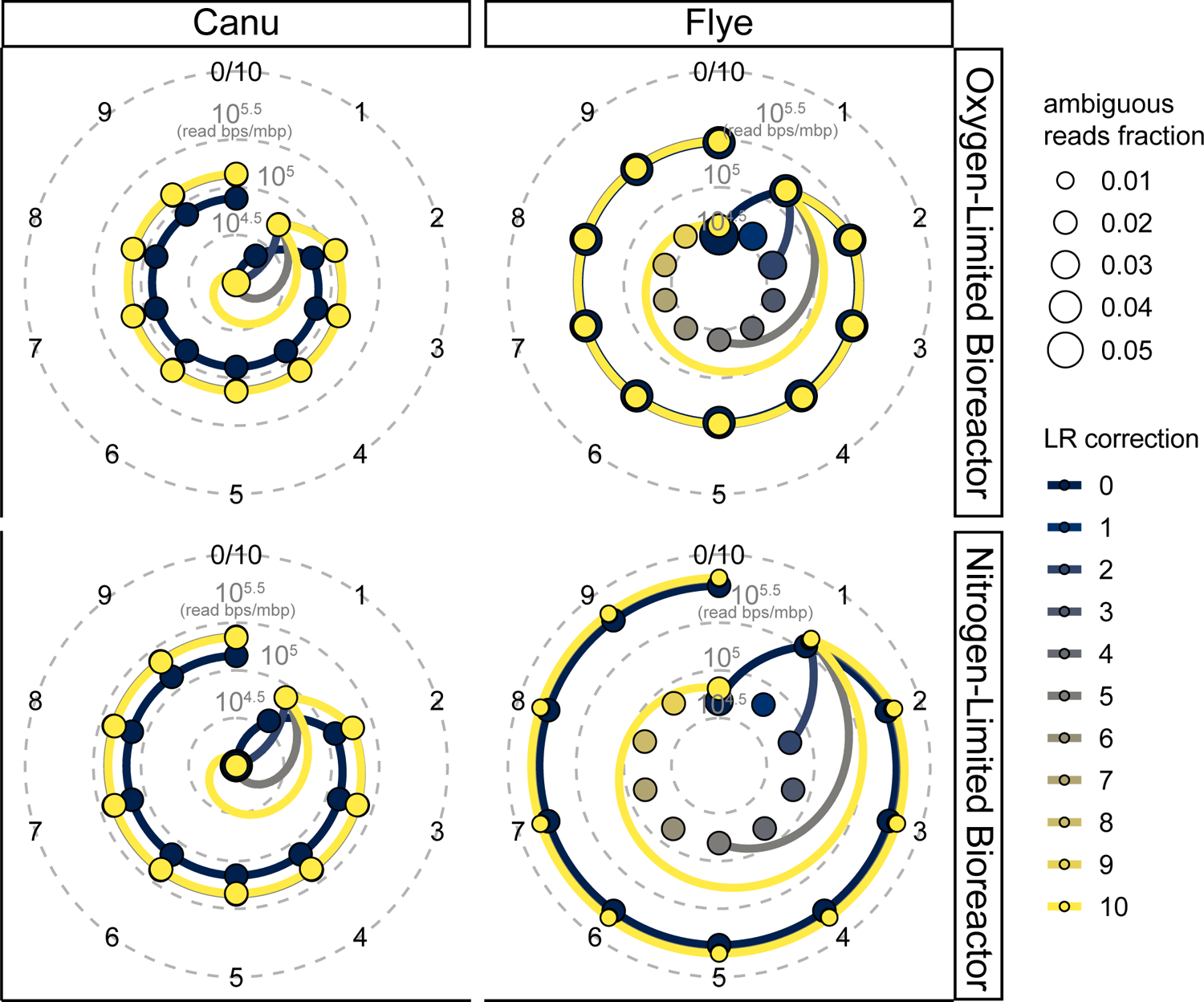
Polar coordinate plot of Short Read (SR) recruitment for each bioreactor and Long Read (LR) assembly throughout the iterative LR correction and SR polishing processes. The two bioreactors are separated over vertical panels, and the two LR assemblers are separated over the horizontal panels. Each panel shows a polar coordinate plot of the LR correction or SR polishing iteration in a circle with the bps in SR recruited per contig per Mbp sequence length as the distance from the center. The innermost ten points (due to lower values) indicate the amount of bps in SR recruited after up to ten LR correction iterations without SR polishing. The colored lines connecting points of the same color link assemblies with the same number of LR correction steps preceding SR polishing. The size of the points indicates the fraction of SR that were ambiguously recruited, i.e., SR aligning to 2 or more positions.

## Discussion

### Correction and polishing affect gene- and genome-centric community reconstruction

Hybrid assembly approaches, leveraging complementary beneficial attributes of both LR and SR platforms to overcome their limitations, are already being used to study microbial communities in various evniromenments^1–15^. Assembly of LRs offers a major advantage over SR data alone, due to the ability to achieve greater contiguity, however, it comes at a significant cost due to lower accuracy^1,5,8,11,14,17,21–24,32^. Each study, from benchmarking to community reconstructions, applied different methods in their hybrid assembly approaches, and it’s probable that the biological and technical complexity of microbial community metagenomes limits the implementation of uniform methods across different studies. One of the least consistently applied, and likely dataset- and tool-dependent, aspects of applying a hybrid assembly approach for a microbial community metagenome is the iterative processes of LR correction and SR polishing to improve the accuracy of LR assemblies.

Few studies have explored the impacts of these iterative processes, despite being implemented by most of them. Generally, the consensus we understood was that while SR polishing offers the greatest improvements, some LR correction is advantageous, and both offer diminishing returns^13,17,21,26,31,37^. We first showed that microbial community composition changes during these iterative processes, which was, as far as we know, the first demonstration of this phenomenon. We further showed that genome recovery was also greatly impacted by these processes, which combined with marker gene beta diversity, demonstrates that the exact number of iterations for these processes has a significant impact on reconstruction, and thus also interpretation, of the microbial communities. Fortunately, results here followed similar patterns with each other, and also in many aspects mirrored published studies comparing different aspects of hybrid assembly across multiple tools and over several iterations^1,2,4,7,10,12–15,17,18,22,26–31,37^. However, there is not only substantial methodological variation between published approaches and the performed here, but also biological variation between each dataset which probably precludes the development and application of a universal or standardized approach for applying hybrid metagenomics to microbial communities. Therefore, it is likely that each dataset needs to be empirically evaluated and its hybrid assembly approach optimized.

### Simple, reference-independent assembly characteristics as proxies for the quality of hybrid assemblies of microbial community metagenomes

Further complicating hybrid assembly for microbial community metagenomes is the lack of a clear assessment of quality. While mis-assembly and contiguity are the most common metrics, these assembly characteristics either require high-quality reference genomes that may not exist for poorly characterized ecosystems, or would not substantially change while optimizing LR correction or SR polishing^5,12,17,18,22,23,26–28,30–33,37,44^. Therefore, assembly characteristics appropriate for a microbial community should be dataset-dependent and expected to change as assembly accuracy also changes.

Here, several assembly characteristics were tracked during the iterative processes to determine which were suitable proxies for assembly quality, and to help optimize hybrid assembly of uncharacterized microbial community metagenomes. Specifically, gene fragmentation and SR recruitment were focused on during LR correction and SR polishing because these features have already been proposed as suitable proxies for hybrid assembly of microbial genomes^4,11–13,17,22,45^. These simple assembly characteristics followed patterns consistent many benchmarking and comparative studies^4,12,13,17,22,26–31:^ (1) the greatest improvements occurred within the first three iterations of SR polishing; (2) LR correction improved the accuracy of the assemblies but was not always apparent until after SR polishing; (3) beyond five iterations LR correction or SR polishing offered severely diminished returns. Furthermore, simpler and reference-independent assembly characteristics were strongly correlated with more advanced, reference-dependent characteristics, *i.e.*, gene counts, lengths, and bps in genes were all correlated with the gene fragmentation profiles, as well as automated bin recovery and marker gene beta diversity trajectories. Thus, these simple, reference-independent assembly characteristics are reliable indicators of assembly accuracy and can be used to empirically optimize hybrid assembly of uncharacterized microbial community metagenomes.

### Unexplored factors and limitations of this study

The approach in this study was not without issue, and therefore here we describe expected deviations from our approach that may be necessary for certain users or datasets. First, recent technological advances for both LR sequencing and application may ease the approach for integrating hybrid metagenomic assembly of microbial communities, but this study was performed using older sequencing chemistry and basecalling algorithms. Some differences remain, but both major LR platforms, Pacific Biosciences (PacBio) and Oxford Nanopore Technologies (ONT) have achieved large improvements in chemistry and basecalling that have increased both depth and accuracy of LR sequencing^4,11,12,24,45,72,73^. Therefore, not only will LR-alone assemblies be more accurate as a result of these advances, they already may be sufficient for retrieving HQ genomes without SR polishing^12,74^. However, resources may restrict users to older and more accessible technologies, or projects may already have older datasets that still need analysis that integrate short and long read technologies. Additionally, for the foreseeable future, SR complements to LR datasets will likely continue to increase the sample count for differential coverage binning. Furthermore, SRs still appear to aid assembly accuracy for high-quality microbial genomes reconstructed from LR^75^. Therefore, combining both sequencing technologies will be attractive as a cost-effective approach yielding the highest quality data compared to any single platform^12^.

Second, the myriad of programs and pipelines to process LRs from basecalling, to read correction, to assembly, to assembly correction, SR recruitment and polishing, and more, continuously improve and expand^3,4,14,17,18,21–24,26,27,29,31,33–38,40–43,45,49,52,54,55,70,71,76–89.^ To reasonably perform this study and analyze the results, only two assembly, one LR correction, and one SR polishing programs were tested, thus introducing technical variation by assembler but limiting it during LR correction and SR polishing. These programs were chosen largely based on their observed prevalence in literature and online resources, but we acknowledge that, in particular, Racon and Pilon may be outcompeted by others that yield better results^17,21,26,31,35,37,41,43,76–80^. The programs we used are also dataset agnostic, and have relatively minimal computational requirements in contrast to tools that are dataset-dependent or demand advanced/specific computational capabilities, making them a relatively universal option. For several available LR or hybrid assembly programs, Racon or Pilon have are already implemented^19,26,29^, indicating their value to the field despite the fact that others or various combinations may yield more accurate assemblies. We further note that each of these computational tools also come with a suite of settings to finetune performance that were not explored. One setting of particular interest that was not investigated here, nor could we find literature on, was read recruitment stringency. Theoretically, this could greatly impact optimization of the iterative processes, and thus accurate reconstructions, of microbial communities with multiple similar strains, genomes with multicopy genes or repeat regions, or other factors that have plagued metagenomic studies since their inception.

Third, contig pools were not separated for some analyses in order to simplify the workflow. Specifically, in this workflow circular contigs that likely constituted nearly complete genomes were not removed from the assemblies prior LR correction or SR polishing. We speculate that this simplification could result in mis-assemblies derived from reads, particularly SR, that should align to different strains ambiguously aligning to conserved regions. However, SR ambiguity decreased during iterative error fixing processes, possibly due to the availability of multiple reference sequences to differentially recruit ambiguous reads during SR polishing.

Fourth and lastly, we acknowledge that the biomass collected for DNA isolation and sequencing with the different platforms were separated by several months. Though largely stable after enrichment for over five years, the communities still shift slowly, thought largely due to strains competing for the same niches that will be examined in follow-up studies. These differences could impact ideal integration the LR and SR data if strains substantially shifted in abundance, leading to improper error-fixing events and could produce inaccurate consensus genomes rather than a realistic strain representation.

## Conclusions

Integrating LR and SR sequencing platforms for hybrid assembly of a microbial community metagenome is challenging due to both biological and technical complexities. While improvements to the quality of LR themselves may be able to preclude the need for SRs, we expect that SRs will still be beneficial as relatively inexpensive additional data for differential coverage binning, and users may still desire to explore SR polishing to maximize their assembly quality. Benchmarks combining these technologies have led to some consensus in the methods for hybrid assembly, but direct replication of the methods may not serve all applications. Here, we’ve demonstrated the that simple, reference-free characteristics, specifically gene lengths and SR recruitment, serve as dataset-dependent indicators of microbial community metagenomic assembly quality, and even approximate advanced reference-dependent characteristics. Additionally, the trajectories of reference-independent assembly characteristics were consistent with benchmarks and published workflows, but did also show that the number of iterations can have substantial impact on downstream analyses. In summary, assembly characteristics that are simple, reference-independent, and precursors to downstream analyses, *e.g.*, gene counts and bps, or recruited short read profiles, can help optimize hybrid assembly approach for microbial community metagenomes. Rather than inform other users of the best approach for their own hybrid metagenomic microbial community projects, we provide others with assembly-level characteristics that are proxies quality to help optimize their own approach. Finally, we offer practical recommendations to enable other users to integrate LR and SR sequencing platforms for microbial community metagenomes with fewer obstacles and guided by empirical data from their own projects.

### Additional practical recommendations

- Use the same pool or replicate extractions of DNA for both SR and LR, which ideally minimizes community differences that might lead to mis-assembly.
- Calculate read depth using bps rather than read counts when mixing LR and SR profiles, otherwise LR depth will be vastly and unevenly underestimated.
- If possible, use multiple assembly programs prior to LR correction and SR polishing, and then consolidate the genome outputs (circular, MQ automated bins) because different algorithms may have a substantial impact on which community members assemble well^22^.
- LR correction should precede SR polishing to avoid the (re-)introduction of errors.
- A mixture of different SR polishing tools may yield the best overall quality^17^, likely due to differential abilities to polish specific errors.
- These analyses are all relative, *i.e.*, dataset-dependent, and therefore should not be used to compare different systems.
- While the ultimate goal might be genome recovery, we suggest monitoring gene fragmentation, read recruitment, or data collected during LR correction or SR polishing for empirically guiding optimization.

`o` Logged data from some programs can be information-rich and should be mined when possible. For example, Pilon records the changes it makes, which also followed the same patterns as the other analyses, *i.e.*, the most and largest changes occur in the first few iterations but plateaued on a log-scale by ∼4 iterations (Fig. S11).

## Supporting information

Suppplemental File 1

