## Supplementary material for "Simple, reference-independent analyses help optimize hybrid assembly of microbial community metagenomes": Suppplemental File 1

####### File_S1
####### Compiled example code
# Note 1: files, directories, and executables have been made generic to simplify the code.
# Note 2: threads and memory allocations have been removed as these are both data- and computing resource-dependent
# Note 3: most processes were looped to use the exact same settings for both bioreactors or both long read assemblies, but for simplicity the code is shown as though there was a single bioreactor and/or long read assembly

##### Read QC
### Long reads, ie Oxford Nanopore Technologies platform
reformat.sh in=ONT.fq out=temprds.fq minlen=3000 # reformat.sh is a BBtools utility
porechop -i temprds.fq -o ONT.qc.fq --min_split_read_size 3000

### Short reads, ie Illumina MiSeq platform
bbduk.sh in1=MS_R1.fq in2=MS_R2.fq out1=MS_R1.qc.fq out2=MS_R2.qc.fq qtrim=rl trimq=18 minlength=200 ktrim=r ref=adapters,artifacts,phix,lambda,mtst

##### Long read subsampling
trycycler subsample -r ONT.qc.fq -o subsampled/ # creates 12 by default in the "subsampled/" directory

##### Short read and short-read first hybrid assembly
### Unicycler, short reads only
python3 unicycler-runner.py -o Unicycler_short --min_fasta_length 1000 -1 MS_R1.qc.fq -2 MS_R2.qc.fq

### Unicycler, long reads only
python3 unicycler-runner.py -o Unicycler_long --min_fasta_length 1000 -l ONT.qc.fq

### Unicycler, hybrid
python3 unicycler-runner.py -o Unicycler_hyb --min_fasta_length 1000 -1 MS_R1.qc.fq -2 MS_R2.qc.fq -l ONT.qc.fq

### SPAdes, short reads only
python3 spades.py --meta -k 27,47,63,77,89,99,107,115,121,127 -o Spades_short -1 MS_R1.qc.fq -2 MS_R2.qc.fq # k-mers match those picked by Unicycler for short read only assembly

### SPAdes, long reads only
python3 spades.py --meta -o Spades_long --nanopore ONT.qc.fq

### SPAdes, hybrid
python3 spades.py --meta -k 27,47,63,77,89,99,107,115,121,127 -o Spades_hyb -1 MS_R1.qc.fq -2 MS_R2.qc.fq --nanopore ONT.qc.fq # k-mers match those picked by Unicycler for short read only assembly

### OPERA-MS, hybrid
perl OPERA-MS.pl --out-dir Opera_hyb --contig-len-thr 1000 --no-ref-clustering --short-read-assembler spades --long-read-mapper minimap2 --short-read1 MS_R1.qc.fq --short-read2 MS_R2.qc.fq --long-read ONT.qc.fq

##### Long read assembly
### Canu, default
canu -p Canu_def_asm -d ONT_canudef_asm/ genomeSize=1m -nanopore-raw ONT.qc.fq

### Canu, metagenome-optimized
canu -p Canu_meta_asm -d ONT_canumeta_asm/ corMinCoverage=0 corOutCoverage=all corMhapSensitivity=high correctedErrorRate=0.105 genomeSize=5m corMaxEvidenceCoverageLocal=10 corMaxEvidenceCoverageGlobal=10 -nanopore-raw ONT.qc.fq

### Canu, metagenome-optimized with subsampled reads
for reads in subsampled/*qc.fq ; do
canu -p Canu_sub_"${reads/.qc.fq/_asm}" -d Canu_sub/Canu_"${reads/.qc.fq/_asm}" corMinCoverage=0 corOutCoverage=all stopOnLowCoverage=0.5 corMhapSensitivity=high correctedErrorRate=0.105 genomeSize=5m corMaxEvidenceCoverageLocal=10 corMaxEvidenceCoverageGlobal=10 -nanopore-raw $reads
done

### Flye, default
flye -o Flye_def_asm/ --nano-raw ONT.qc.fq

### Flye, metagenome-optimized
flye -o Flye_meta_asm --nano-raw ONT.qc.fq --meta

### Flye, metagenome-optimized with subsampled reads
for reads in subsampled/*qc.fq ; do
flye -o Flye_sub/Flye_sub_"${reads/.qc.fq/_asm}" --nano-raw $reads --meta
done

##### Compile initial assemblies into one location
mkdir asms_init
for asm in */*_asm* ; do
ASM1=${asm/*\//}
ASM=${ASM1/.fasta/.fa} # this assumes that the output assembly names are relatively similar from all programs with the exception of ".fasta" vs ".fa" , but the truth is that the outputs are not easily generalized and more complicated code (e.g. per assembler) is necessary to make the names harmonious
reformat.sh in="$asm" out=asms_init/"$ASM" minlen=4000 # reformat.sh is a BBtools utility
done

##### Subset circular and long (>1 Mbp) contigs, then estimate quality and infer phylogeny
mkdir asms_init_subsets/{circ,long,checkm,gtdb}
mkdir -p asms_init_subsets/circ/indivseqs/
mkdir -p asms_init_subsets/long/indivseqs/
for asm in asms_init/*fa
ASM1=${asm/*\//}
ASM=${ASM1/_asm.fa/_long.fa}
# Note, there is no uniform way that assembly programs output circularity information, so here it has been assumed that for each assembly the user has identified the circular contigs and put them all in one list named "circularcontigs.list"
filterbyname.sh in="$asm" out=asms_init_subsets/circ/"$ASM"_circ.fa names=circularcontigs.list keep=t # filterbyname.sh is a BBtools utility
filterbyname.sh in="$asm" out=asms_init_subsets/long/"$ASM"_long.fa names=circularcontigs.list minlen=1000000 keep=f # filterbyname.sh is a BBtools utility
grep '>' asms_init_subsets/circ/"$ASM"_circ.fa | sed 's|>||' asms_init_subsets/circ/"$ASM"_circ.list
	for seq in $(< asms_init_subsets/circ/"$ASM"_circ.list ) ; do
	filterbyname.sh in=asms_init_subsets/circ/"$ASM"_circ.fa out=asms_init_subsets/circ/indivseqs/"$ASM"_"$seq".fa name="$seq" keep=t # filterbyname.sh is a BBtools utility
	done
grep '>' asms_init_subsets/long/"$ASM"_long.fa | sed 's|>||' asms_init_subsets/long/"$ASM"_long.list
	for seq in $(< asms_init_subsets/long/"$ASM"_long.list ) ; do
	filterbyname.sh in=asms_init_subsets/long/"$ASM"_long.fa out=asms_init_subsets/long/indivseqs/"$ASM"_"$seq".fa name="$seq" keep=t # filterbyname.sh is a BBtools utility
	done
done
for subset in circ long ; do 
checkm lineage_wf -f asms_init_subsets/checkm/"$subset"/"$subset"_checkm.txt --tab_table -x fa asms_init_subsets/"$subset"/indivseqs/ asms_init_subsets/checkm/"$subset"
gtdbtk classify_wf --genome_dir asms_init_subsets/"$subset"/indivseqs/" --out_dir asms_init_subsets/gtdb/"$subset"_gtdb -x fa

##### Long read correction of long read assembly
mkdir lr_corrected
for lrasm in asms_init/*_meta_asm.fa ; do 
asmblr1=${lrasm/asms_init\//}
asmblr=${asmblr1/_meta*/}
minimap2 -c -x map-ont $lrasm ONT.qc.fq > ONT_map_"$asmblr".paf
racon  -u ONT.qc.fq ONT_map_"$asmblr".paf $lrasm > lr_corrected/"$asmblr"_rac1_asm.fa
	for n in $( seq 1 9 ) ; do
	newn=$[ n + 1 ]
	minimap2 -c -x map-ont lr_corrected/"$asmblr"_rac"$n"_asm.fa ONT.qc.fq > ONT_map_"$asmblr"_rac"$n".paf
	racon -u ONT.qc.fq ONT_map_"$asmblr"_rac"$n".paf lr_corrected/"$asmblr"_rac"$n"_asm.fa >lr_corrected/"$asmblr"_rac"$newn"_asm.fa
	done
done

##### Short read polishing of long read assembly
mkdir sr_polished
for lrasm in asms_init/*meta_asm.fa lrcorrected/*rac2* lrcorrected/*rac5* lrcorrected/*rac10* ; do 
asm1=${lrasm/*\//}
asm2=${asm1/_meta/}
asm=${asm2/_asm.fa/}
bbmap.sh ref="$lrasm" in1=MS_R1.qc.fq in2=MS_R2.qc.fq outm=MS_map_"$asm".bam minid=0.9 idfilter=0.975 ambiguous=all pairedonly=t bs=bs.sh nodisk; sh bs.sh
pilon --genome $lrasm --bam MS_map_"$asm"_sorted.bam --output "$asm"_pil1_asm.fa --outdir sr_polished --changes --fix all --duplicates 
	for n in $( seq 1 9 ) ; do
	newn=$[ n + 1 ]
	bbmap.sh ref=sr_polished/"$asm"_pil"$n"_asm.fa in1=MS_R1.qc.fq in2=MS_R2.qc.fq outm=MS_map_"$asm"_pil"$n".bam minid=0.9 idfilter=0.975 ambiguous=all pairedonly=t bs=bs.sh nodisk ; sh bs.sh
	pilon --genome sr_polished/"$asm"_pil"$n"_asm.fa --bam MS_map_"$asm"_pil"$n"_sorted.bam --output "$asm"_pil"$newn"_asm.fa --outdir sr_polished --changes --fix all --duplicates
	done
done

##### Compile all assemblies into one location
mkdir ALL_ASMS
cp asms_init/*fa ALL_ASMS/
cp lr_corrected/*fa ALL_ASMS/
cp sr_polished/*fa ALL_ASMS/

##### Assembly quality estimation
### General statistics using Quast
metaquast.py -o mQuast --max-ref-number 0 ALL_ASMS/*fa

### Open reading frame identification using Prodigal
mkdir prodigal/{GBK,FAA,FNA}
for asm in ALL_ASMS/*fa
ASM1=${asm/ALL_ASMS\//}
ASM=${ASM1/_asm.fa}
prodigal -i $asm -o prodigal/GBK/"$ASM".gbk -a prodigal/FAA/"$ASM".faa -d prodigal/FNA/"$ASM".fna -p meta
sed 's| |.*|' prodigal/FAA/"$ASM".faa > prodigal/FAA/"$ASM"_cln.faa 
sed 's| |.*|' prodigal/FNA/"$ASM".fna > prodigal/FNA/"$ASM"_cln.fna 
done

### Gene fragmentation estimation using IDEEL
mkdir ideel/{genomes,proteins} # IDEEL is by default set up to run in its install path, so either adjust IDEEL's executables or copy data to the install location to run. 
cp ALL_ASMS/*fa ideel/genomes/
cp prodigal/FAA/*faa ideel/proteins/
cd ideel
snakemake --use-conda all

### Assembly-level marker gene fragmention and redundancy using BUSCO
mkdir busco/
for faa in prodigal/FAA/*faa
FAA1=${faa/*\//}
FAA=${FAA1/.faa}
busco -i $faa -o $FAA --out_path busco/ -m prot -l bacteria_odb10 --download_path busco/
done

### Short read recruitment using BBMap
mkdir bbmapstats/
for asm in ALL_ASMS/*fa
ASM1=${asm/ALL_ASMS\//}
ASM=${ASM1/_asm.fa}
bbmap.sh ref="$asm" in1=MS_R1.qc.fq in2=MS_R2.qc.fq perfectmode=t ambiguous=random mappedonly=t pairedonly=t statsfile=bbmapstats/"$ASM"_stats.txt scafstats=bbmapstats/"$ASM"_scaf.txt rpkm=bbmapstats/"$ASM"_rpkm.txt covstats=bbmapstats/"$ASM"_cov.txt nodisk
done

### Assembly-level marker gene redundancy using CheckM
checkm taxonomy_wf -f checkm/checkm.txt --tab_table -x faa domain Bacteria prodigal/FAA/ checkm/

### Metabat2, automated binning
cat MS_R1.qc.fq MS_R2.qc.fq > MSreads # precursor for mmlong utility
mkdir mbat/{mapping,allbins,checkm,busco}
for asm in ALL_ASMS/*fa
ASM1=${asm/ALL_ASMS\//}
ASM=${ASM1/_asm.fa}
bash /mmlong/mmlong_tools/mmlong_readcoverage2 -a $asm -d MSreads -m ilm -o mbat/mapping 
sed "s|coverage|MS_cov|" mbat/mapping/MSreads_cov.csv | sed "s|stdev|MS_var|" > mbat/mapping/MS_map_"$ASM"_cov.csv
bash /mmlong/mmlong_tools/mmlong_readcoverage2 -a $asm -d ONT.qc.fq -m np -o mbat/mapping
sed "s|coverage|ONT_cov|" mbat/mapping/ONT.qc.fq_cov.csv | sed "s|stdev|ONT_var|" > mbat/mapping/ONT_map_"$ASM"_cov.csv
grep '>' $asm | sed 's|>||' > contigs
R $ASM
library("tidyverse")
args <- commandArgs() 
l1=list.files(pattern="contigs")
l2=sort(list.files(pattern=paste0("mbat/mapping/.*map_",args[N],"_cov.csv"))) # adjust the args[N] to call the variable from the command line suit R setup
out=c(l1,l2) %>% 
	lapply(read_csv) %>% 
	reduce(.,full_join,by="scaffold") # works in a test... not sure why or how... or why or how I cannot do it another way
write_tsv(out,paste0("mbat/mapping/",args[N],"_depth.tsv") # adjust the args[N] to call the variable from the command line suit R setup
q()
metabat2 -i $asm -a mbat/mapping/"$ASM"_depth.tsv -o mbat/"$ASM"/"$ASM" --saveCls --unbinned
cp mbat/"$ASM"/"$ASM".[0-9][0-9]*.fa mbat/allbins/
done
checkm taxonomy_wf -f mbat/checkm/checkm.txt --tab_table -x fa domain Bacteria mbat/allbins/ mbat/checkm
for bin in mbat/allbins/*fa ; do
BIN1=${bin/*\//}
BIN=${BIN1/.fa/}
busco -i $bin -o "$BIN" --out_path mbat/busco/ -m genome -l bacteria_odb10 --download_path mbat/busco/
done
seqkit stats mbat/allbins/*fa | sed 's|  *|\t|' > mbat/seqkitstats.tsv

### Identification and depth quantification of rpoB genes using HMMscan, BBtools utilities, and mmlong utilities
cat MS_R1.qc.fq MS_R2.qc.fq > MSreads # precursor for mmlong utility
mkdir rpob/{tbls,contigs,genes,proteins,mapping}
cp /HMMdbs/PF04563.hm* rpob/ # HMM downloaded and prepared 2020-06-09
for faa in prodigal/FAA/*_cln.faa 
FAA1=${faa/*\//}
FAA=${FAA1/_cln.faa}
ASM=${FAA1/_cln.faa/_asm.fa}
FNA1=${faa/FAA/FNA}
FNA=${FNA1/faa/fna}
hmmscan --seed 42 -Z 47079205 -E 1000 --tblout rpob/tbls/"$FAA"_rpob_seq.txt -o rpob/tbls/"$FAA"_rpob.txt rpob/PF04563.hmm $faa 
grep 'PF04563.15 ' rpob/tbls/"$FAA"_rpob_seq.txt | sed 's|.*PF04563.15 ||' | sed 's|  *|	|g' | awk -F '\t' '{ if ( $3 <= 1e-3  && $4 >= 20 ) print $1 }' > genes.list
sed 's|_[0-9][0-9]* ||' genes.list > contigs.list 
filterbyname.sh in="$faa" out=rpob/proteins/"$FAA" names=genes.list include=t # filterbyname.sh is a BBtools utility
filterbyname.sh in="$fna" out=rpob/genes/"$FNA" names=genes.list include=t # filterbyname.sh is a BBtools utility
filterbyname.sh in=ALL_ASMS/"$ASM" out=rpob/contigs/"$ASM".fa names=contigs.list include=t # filterbyname.sh is a BBtools utility
done
for rpocont in rpob/contigs/*fa
rpoCONT1=${rpocont/*\//}
rpoCONT=${rpoCONT1/.fa/}
bash /mmlong/mmlong_tools/mmlong_readcoverage2 -a $rpocont -d MSreads -m ilm -o rpob/mapping 
sed "s|coverage|MS_cov|" rpob/mapping/MSreads_cov.csv | sed "s|stdev|MS_var|" > rpob/mapping/MS_map_"$rpoCONT"_cov.csv
bash /mmlong/mmlong_tools/mmlong_readcoverage2 -a $rpocont -d ONT.qc.fq -m np -o rpob/mapping
sed "s|coverage|ONT_cov|" rpob/mapping/ONT.qc.fq_cov.csv | sed "s|stdev|ONT_var|" > rpob/mapping/ONT_map_"$rpoCONT"_cov.csv
grep '>' $rpocont | sed 's|>||' > contigs
R $ASM
library("tidyverse")
args <- commandArgs() 
l1=list.files(pattern="contigs")
l2=sort(list.files(pattern=paste0("rpob/mapping/.*map_",args[N],"_cov.csv"))) # adjust the args[N] to call the variable from the command line suit R setup
out=c(l1,l2) %>% 
	lapply(read_csv) %>% 
	reduce(.,full_join,by="scaffold") # works in a test... not sure why or how... or why or how I cannot do it another way
write_tsv(out,paste0("rpob/mapping/",args[N],"_depth.tsv") # adjust the args[N] to call the variable from the command line suit R setup
q()
