## Supplemental Text 1 for "Simple, reference-independent analyses help optimize hybrid assembly of microbial community metagenomes"

**Title:**

**Affiliations:**

### Supplemental Methods

*Approach justification*

Benchmarking has shown that every tool choice, version, and implementation – the sequencing platform, basecalling, assembly, correction and polishing, and more – will affect (meta)genome reconstruction quality due to differences in chemistry, algorithms, or other features[1–68]. There does not appear to be any single or combination of technologies that yields a perfect assembly, and a “best” option is subjective, for example, given willingness to sacrifice computational demands for weaker performance, or vice-versa. The assembly quality metric(s), or even the tools used to assess them, can arguably also impact the true quality of the reconstructed (meta)genome as well, particularly for genome recovery[69–73]. The approach here was informed by benchmarking or comparative studies, largely of various microbial isolates and synthetic or mock communities, as well as studies of various ecosystems including wastewater treatment sludge that could be considered relatively similar to our bioreactors inoculated with activated sludge[15,16,24,25,27,74]. No quantification of use in the literature was performed, but in general, we chose tools and parameters based on: (1) the prevalence of their use in the field of microbial (meta)genomics, (2) minimizing computational and monetary resource expenditure, which are likely limiting factors for many projects, (3) value inferred from published benchmarking or comparative studies, and (4) ease of use and documentation or incorporation into already existing pipelines/workflows. We did not include several other tools that did meet these criteria, were even tested, or even reportedly performed better to reduce experimental complexity and technical variation, or more simply because they were not available or functional on our computing system without substantial effort.

Biomass collection dates from the bioreactor systems examined here were chosen to update the metagenomic inventory available for the system in preparation for a community reconstruction (Nov. 2020) and then later for the intent of long read sequencing (Mar. 2021).

Two DNA isolation protocols were used because it is standard in our group due to strong evidence that it introduces sufficient bias to aid differential coverage binning[75–77].

The ONT platform was used to generate LR sequencing for these bioreactors because it was an in-house facility capability. Following, only one version of chemistry was available in the facility at the time, and what appeared to be the state-of-the-art basecalling algorithm for microbial genomes was used for all of the facility's outputs[23,78]. Similarly, Illumina MiSeq was used because it was an in-house SR sequencing capability. Both LR and SR sequencing was performed according the standard operating procedures, including reagents and protocols, of the sequencing facility, which were informed by manufacturer instructions. One bioreactor was sequenced more deeply because of an observed greater richness in a 16S rRNA gene amplicon sequencing survey (unpublished), use of this bioreactor for previous metagenomic projects[79] and greater interest in improving upon extant community reconstructions.

BBduk was used to trim and filter SRs because of its one-step capabilities for Illumina read quality processing and its accompanying suite of tools for working with various sequencing data[80]. Porechop was used for LR trimming because of its ease of use and implementation in studies of related ecosystems[16,74,81]. The exact quality cutoffs were arbitrarily chosen to help balance data quantity and quality (not shown).

OPERA-MS appeared to be a prevalent SR-first hybrid assembler but is methodologically biased towards model organisms and host ecosystems[27], whereas hybridSPAdes appeared approximately as prevalent an SR-first hybrid assembler but lacks any methodological bias towards certain taxa or ecosystems and is capable of SR-alone assembly[20]. Unicycler has the ability to perform SR-first hybrid as well as SR- and LR-alone assemblies, which therefore enables assembly of all possible read types and combinations with one program[82]. Additionally, its ability to automate the optimization of the assemblies for successful, high-quality, circular single genome recovery has made it popular. Furthermore, all tested SR-dependent assemblers tested implement SPAdes for SR-assembly, thus reducing the technical variation.

The Trycycler utility "subsample" was used to evenly subsample the LRs for assembly because of its shared developer with Unicycler[8,82] and seeming ease of use. Published comparisons tend to suggest that these assemblers are capable of microbial genome recovery on par with comparable tools[7,8,15,20,24,27,82], and have even pointed out their prevalence in the field[15].

Canu and Flye were used for LR assembly because of the popularity in the field, and publications describing metagenome-optimization or a suite of parameters to enable this aspect[14,15,21,28,74]. As some of the first LR assemblers, these are compared often and tend to compete well for microbial genomes, or are included in comparisons of other tools[6–11,14,15,21,24,32,40–43]. Other assemblers were tested but were not described here because (1) they did not appear to perform as well, (2) were not as prevalent in the field, and therefore (3) would only have added unnecessary additional complexity with potentially little impact on the results. Several settings were tested as well. The metagenome-optimized settings were favored because the goal was to assemble metagenomic data, but default settings were included as few direct comparisons were made and we wanted to ensure advantage over the default for our dataset. Subsampled reads were also tested because Trycycler documentation suggests that slightly different read pools may be assembled slightly differently. There was initially an intent to expand correction and polishing to consensus MAGs from Trycycler, but this was deemed overly complex and consumed unnecessary computational resources. This information was kept in this manuscript because it both validated and contrasted the expectations of benchmarks: slightly different assemblies were generated in terms of size and composition but there wasn't a major apparent sacrifice in quality, in a somewhat assembler-dependent manner.

Racon and Pilon were chosen for LR correction and SR polishing, respectively. In many benchmarking studies, these tools may be outcompeted by others, or may need additional tools to resolve leftover errors[1,8,12,73]. However, these tools are commonly used in the field, are compared frequently and appear to perform nearly on par with other tools, have low computational demands, and even are incorporated into published pipelines or workflows[1,8,14,16,21,22,37,39,43,68,74,82–85]. One tool using Racon in its automated workflow, Unicyclyer[82], was already used here thus limiting unnecessary technical variation. Additionally, these tools have relatively simple computational requirements that allow almost any user to use them. Furthermore, these specific tools have additional advantages: Racon can be used for LR correction or SR polishing, though this complicates a general inference of its success[9,16,74], and Pilon includes several other features like data-rich logging and variant calling[13].

The maximum number of iterations of LR correction and SR polishing, 10, was chosen because it was the maximum observed in the literature[1]. The breaks in LR correction at which SR polishing was performed, 0, 2, 5, and 10, were chosen to represent the SR polishing alone (0), a seemingly common number of iterations (2[6,9,11,16,21,28,32,68,74,84,86–88]), and excessive (5[8,86], 10[1,86,89]) by which we expected to observe clearly diminishing returns on the efforts given information from benchmarks. Highly diverse correction and polishing strategies complicates the ability to inform these arbitrary decisions. We further considered that many benchmarks were performed on single microbial genomes, particularly those that compared assembly quality over multiple iterations of tool application, and the complexity of a metagenome may warrant additional iterations. We did not examine SR polishing after a single iteration of LR correction because it was rarely observed in literature and essentially never when these tools were used.

metaQUAST was used for cursory comparison of assembly statistics because it is commonly employed and provides valuable summaries for comparing contig and bp distributions of multiple assemblies[4,7,10,14,16,19,20,26,37,39,69,82,85,88].

The read recruitment software was chosen for separate reasons. Minimap was chosen for LR recruitment because the output format is recognized by Racon and is implemented in other tools used here, but we acknowledge it's performance may not be competitive with others in some cases though its low resource demand may offset its disadvantages[9,54,61,62,90,91]. BBmap was used for SR recruitment because it offers many thresholds to apply towards alignment parameters and data-rich logging and summaries that support an analyses of the read alignments themselves, and performs on par with comparable tools though it can be computationally demanding in some cases[54,80]. To recruit reads for input to Pilon, a 97.5% identity threshold was applied with to help balance maximizing SR alignment to inaccurate sequences and minimize multiple SR alignments, though no additional thresholds were tested or analyses done to validate this decision. For assessing read recruitment and ambiguity, BBmap was implemented to only report perfect mappings to reduce noise of multiple SR alignments caused by strain diversity and multi-copy genes[80]. The mmlong utility *readcoverage2* was used for abundance profiling because it calculated read depths similarly for both LR and SR datasets (*i.e.*, by bps rather than read counts) and offered a simple output that can be input into the automated binner chosen[91].

Prodigal was chosen for ORF finding because it includes metagenome-optimized settings, information-rich headers that supported some of these analyses, and is implemented in the marker gene recovery pipelines chosen therefore reducing unnecessary technical variation[71,72,92–94].

As far as we know, there are no tools specifically developed to inventory phylogenetic/single copy marker genes to assess the quality, redundancy, or fragmentation of a metagenomic assembly. CheckM was chosen for microbial marker gene recovery summarization and automated bin quality estimation because it is commonly applied for these purposes in microbial genome recovery from metagenomes, is used in benchmarking studies, and has a straightforward implementation in downstream tools[14,16,17,19,24,25,27,28,70,71,74,83,84,88,94]. BUSCO was chosen as a marker gene recovery summary tool partially because, in contrast to CheckM, it is capable of assessing marker gene of Eukaryotes and therefore can be applied to various uncharacterized communities, and is similar to CheckM in its application in the field and benchmarks[1,10,19,28,39,93,95]. An additional advantage of CheckM is a theoretically infinite score for whole-genome redundancy, while an additional advantage of BUSCO is that it assesses the fragmentation of marker genes in addition to their presence.

IDEEL was chosen to compare gene lengths to references sequences because it has been used in similar studies, and has little competition[8,16,84,88].

GTDB-tk was chosen for taxonomic inference of the microbial genomes for essentially the same reasons as CheckM, and also has few competitors[4,8,16,24,25,28,72–74,83–85,88,94–96].

Metabat2 was chosen as the automated binner because it tends to perform well in benchmarks, is common to the field and often compared, and is incorporated into published pipelines[4,16,17,19,24,25,28,50,52,70,74,83–85,88,95,96]. Only one binning program was chosen to reduce technical variation and complexity, and theoretically, with the advantages in contiguity that LR assembly offers for automated binning, we thought that the automated binner(s) should not make a large difference.

The *rpoB* gene was chosen as the single-copy phylogenetic marker because the HMM profile (PF04563) curated by the Protein Family (pfam) database includes all three domains of life and the computational capabilities were in place from a published project for this system[79,97]. We acknowledge that other phylogenetic markers may be better and/or more common.

For beta diversity clustering, we chose NMDS calculated with Bray-Curtis dissimilarity as this is a staple analysis of both macro- and microbial ecology[98–100]. Other statistics, for example, PCA or PCoA, are also popular, but this approach was developed specifically for compositional analyses of communities and performs equally well[98,99,101]. The “species” scores were analyzed instead of the typical “site” scores because the data were structured such that correction and polishing iterations were encoded by the marker genes themselves rather than the read profiles.

The tools used here, their purpose in this study, and the rationale for their use are summarized in Table S1.

**Table S1.** Tool choice, its purpose in this manuscript, and the rationale for or advantages of its use here. Rationale/Advantages in this used to support or justify application in this study over others were almost entirely arbitrary or based on subjective criteria; better-performing tools may exist for the same purpose but were not used for this study. Citations were only provided here for demonstration of specific attributes related to the rationale beyond the tool’s release publication.

| Tool | Purpose | Rationale/Advantages |
|---|---|---|
| Illumina MiSeq (Nextera XT kit) | Short read sequencing | Prevalence in field, in-house capability |
| Oxford Nanopore Technologies MinION (R9.4.1 flowcell) | Long read sequencing | Prevalence in field, in-house capability |

| | | |
|---|---|---|
| Guppy[78] (v 4.0.11, dna_r9.4.1_450bps_hac) | Long read base-calling | Prevalence in field, in-house capability standard |
| BBduk[80] (BBtools v 37.76) | Short read QC | One-step trimming and filtering, companion tools |
| porechop[81] (v 0.2.3_seqan2.1.1) | Long read QC | Prevalence in field |
| hybridSPAdes[20] metaSPAdes[18] (v 3.15.4) | Short read, long read, and short read first hybrid assembly | Prevalence in field, other tools use it therefore reducing technical variation |
| OPERA-MS[27] (v 0.9.0) | Hybrid assembly | Prevalence in field, SPAdes-dependent |
| Unicycler[82] (v 0.4.9b) | Short read, long read, and short read first hybrid assembly | Developed for hybrid assembly to recover high-quality and circular microbial genomes, SPAdes-dependent short read assemblies, long read hybrid assembly using Racon |
| Canu[21] (v 1.8) | Long read assembly | Prevalence in field, often compared (or used in comparative studies), metagenome-optimization possible |
| Flye[14] (v 2.9-b1768) | Long read assembly | Prevalence in field, often compared (or used in comparative studies), metagenome-optimization integrated |
| Trycycler[8] (v 0.4.1) | Long read subsampling | Ease of use |

| | | |
|---|---|---|
| Racon[9] (v 1.3.1) | Long read correction | Prevalence in field, often compared (or used in comparative studies), minimal computational requirements, other tools use it therefore reducing technical variation |
| Pilon[13] (v 1.23) | Short read polishing | Prevalence in field, often compared (or used in comparative studies), minimal computational requirements, data rich logging, companion tools |
| Minimap2[90] (v 2.17-r941) | Long read recruit for correction | Prevalence in field, ease of use with Racon |
| BBmap[80] (BBtools v 37.76) | Short read recruitment for polishing | One-step mapping and filtration, companion tools |
| metQuast[69] (v 5.0.2) | General assembly characteristics | Prevalence in field, ease of use |
| BBmap[80] (BBtools v 37.76) | Short read recruitment analysis | One-step mapping and filtration, data rich output summaries |
| Prodigal[92] (v 2.6.3) | Gene calling for several analyses | Prevalence in field, other tools use it therefore reducing technical variation and reducing computational costs |
| BUSCO[93] (v 5.1.2) | Marker gene identification | Used in field but can be extended more broadly, assesses gene fragmentation |
| CheckM[71] (v 1.1.3) | Marker gene identification | Prevalence in field, assesses gene redundancy |

| | | |
|---|---|---|
| | and microbial genome completion estimation | |
| IDEEL[84] (downloaded 2023-05-13) | Gene fragmentation | Use in field, ease of use, little competition |
| GTDB-tk[72] (v 1.6.0, db v r202) | Microbial taxonomy inference | Prevalence in field, little competition |
| readcoverage2 (mmlong[91] v 0.1.2) | Read depth profiling | Ease of integration with binner, calculated long read depth properly |
| Metabat2[17] (v2.12.1) | Automated binning | Prevalence in field, often compared (or used in comparative studies), low computational requirements |
| RNA polymerase subunit B (*rpoB*) gene (pfam[97] PF04563.15, downloaded 2020-06-09) | Beta diversity | Universal to all three domains of life in essentially single copy, use in field, previous application to the system |
| Non-Metric multi-Dimensional Scaling (NMDS) (vegan[102] v 2.5-7, R[103] v 4.1.2) | Beta diversity | Use in (related) field(s) |

### Supplemental Results

*Additional information on sequencing yield*

**Table S2.** Sequencing yields for both reactors and sequencing platforms after quality control and trimming. OLR, oxygen-limited bioreactor; NLR, nitrogen-limited bioreactor; MiSeq, Illumina MiSeq platform; ONT, Oxford Nanopore Technologies platform.

| Reactor | Sequencer | Read count* | Read bps (total) | Read length (mean) |
|---|---|---|---|---|
| OLR | MiSeq | 7,568,591 | 4,063,854,882 | 268 |
| OLR | ONT | 318,070 | 3,128,778,886 | 9,837 |
| NLR | MiSeq | 8,733,498 | 4,601,661,593 | 263 |
| NLR | ONT | 416,527 | 4,014,258,886 | 9,638 |

* Paired read counts for Illumina data.

*Additional information on the long read, short read, and short read-first hybrid assembly baselines*

SR-alone and SR-first assemblies with SPAdes led to the most fractured assemblies with only 25-37% of the 95-325 total assembled Mbps on larger contigs (>50 Kbp) and did not produce any circularized contigs (Fig. S1). OPERA-MS, which depends on SPAdes in its SR-first hybrid assembly strategy, only improved upon the SPAdes SR-first hybrid assembly by having a larger fraction of bps on larger contigs, almost 50% of the 144-228 total assembled Mbp. All assemblies produced with Unicycler were smaller but more contiguous than the other SR hybrid assembly programs, with 50-90% of the total 57-102 Mbps in larger contigs (>50Kbp) and 19-49 circular contigs. Given that Unicycler was developed for single genome assemblies, these assemblies were expected to be smaller but more contiguous, and likely of higher quality, when, arguably incorrectly, applied to a metagenome. Overall, these programs performed as expected for SR-alone or SR-first hybrid assemblers that can create large assemblies, which tend to be fractured among small contigs and challenging to circularize.

Both LR assemblers Canu and Flye resulted in larger and more contiguous assemblies than the SR assemblers. LR assemblers averaged 170 and 235 total assembled Mbps for the OLR and NLR, respectively, with 50-90% among larger contigs. As expected, assemblies generated with default settings developed for assembly of individual and often Eukaryotic genomes were relatively poor quality compared to metagenome-optimized settings in either size or contiguity. Specifically, the size of the Canu assemblies increased nearly 2-fold by using metagenome-optimized settings, the fraction on larger contigs increased from 52% to 62-66%, and the total bps in circular contigs increased two to three orders of magnitude. Flye's improvements were smaller by increasing total size by 10-20 Mbps and total bps in circular contigs by less than one order of magnitude. Assembling sub-sampled reads slightly decreased assembly sizes and typically maintained contiguity, but interestingly produced the longest contiguous sequence (9.19 Mbps in the OLR) and was able to circularize microbial genomes that were not circularized in the full dataset, including the largest circular contigs (7.17 Mbp in the NLR, see below). Most studies on the effect of read depth demonstrate that increasing depth leads to improved assembly quality, therefore it was unexpected to find that lower read depths helped to recover distinct, presumably complete genomes. Unexpectedly, the correct-then-assemble program Canu yielded smaller and more fractured assemblies compared to the assemble-then-correct program Flye, with 15-37 less total assembled Mbps and only 62-66% versus 88-90% of bps on larger contigs.

Not only did Canu lead to smaller and more fractured assemblies, but, these assemblies appeared to be of lower quality compared to Flye as well. Canu assembled fewer and smaller circular contigs that may represent complete plasmid, phage, or microbial sequences (Fig. S2). Using default settings, it assembled only 3 total circular contigs, none of which were probable microbial genomes (>1 Mbp), while Flye constructed a total of 81 total circular contigs of which 6 and 2 in the OLR and NLR, respectively, may represent complete microbial genomes. This gap hardly narrowed when using metagenome-optimized settings or with sub-sampled reads, resulting in Flye circularizing 13.2-fold more contigs across all assemblies than Canu. Furthermore, the estimated completion of circular contigs from Canu tended to be lower. For circular contigs >5 Kbp and with completion >0%, the mean completion from Canu assemblies was 40.7% across both reactors and 56.5% for Flye. We do acknowledge that Unicycler produced the highest quality circular contigs, with 3 >95% complete and <5% contaminated contigs from SR-first hybrid assemblies, but the overall yield of circularized, larger, and at least partially complete circular contigs was outmatched by both Canu and Flye when run with metagenome-optimized settings. Therefore, when LRs are available, using an LR assembler yields more and more contiguous data but requires error-fixing to match SR-first hybrid assemblies.

Similar trends were observed for non-circular, long contigs (>1 Mbp) that may also represent nearly or essentially complete microbial genomes (Fig. S3). SR-dependent assemblers as well as Canu with default settings tended to produce less than 10 of these larger contigs, while Flye with default and both LR assemblers with metagenome-optimized settings and sub-sampled reads yielded 10-21. Again, we acknowledge that SR-first hybrid assemblers produced the highest-quality long contigs, with 4 contigs from Unicycler and 2 from OPERA-MS representing nearly complete bacterial genomes with >90% completion and <5% contamination. Still, LR assemblers produced more than twice the number of these contigs. Additionally, there were several contigs assembled by Canu and Flye that had unusually large estimated contamination values (12-74%), primarily in sub-sampled read assemblies and Canu assemblies. While it is possible that marker gene redundancy varies greatly by lineage and reaches >10% in isolates, it is also possible that these may be erroneous chimeric contigs composed of at least two genomes, or caused by small-scale errors that lead to gene fragmentation known to affect LRs and their assemblies. In contrast to the circular contigs, there was a larger difference in the estimated completion of these long contigs between bioreactors rather than between LR assemblers – a mean completion of 17.9% for contigs with bacterial phylogenetic markers in the OLR compared to 28.6% in the NLR. The number of contigs matching this criterion was nearly 1.7-fold greater in the NLR than OLR, a phenomenon that we infer to be an artifact of the greater read depth for the NLR.

In summary, the assembly strategy has a profound effect on the outcome of the initial assemblies, from overall size to contiguity and quality, as expected from most benchmarking studies. SR-first hybrid assemblies immediately yield high-quality microbial genomic data, but only for a few members of the community while the rest remain fragmented. On the other hand, LR assemblies produce larger and more contiguous contigs, which require further processing to yield high-quality microbial genomic data. The benefit of these LR assemblies is that given their size and contiguity, they can theoretically enable the reconstruction of larger portions of the microbial communities. The difference between SR and LR assemblies in potential nearly complete microbial genomic information was most apparent when comparing the number of larger circular contigs.

*Additional information on microbial community beta diversity*

On average, 21.5 *rpoB* genes were assembled initially, though there were clear differences in both bioreactor and assembler – the NLR assemblies contained more *rpoB* genes indicating a more diverse system, and Flye led to the recovery of more *rpoB* genes, thus better capturing the communities within both systems (Fig. S4). The average *rpoB* gene recovery increased to 24.2 with LR correction alone, but to at least 25 after just one SR polishing iteration without LR correction. Besides increasing in count, the average length of these genes increased 3.3-fold after SR polishing alone, 3.4-fold after both LR correction and SR polishing, but only 1.2-fold after LR correction alone. Intriguingly, the Canu assembly of the NLR had on average 1.2-fold longer *rpoB* genes after SR polishing compared to the Flye assembly, suggesting that the choice of assembler may have a major impact even ongene-centric community-level analyses. We note that there were on average 8 more *rpoB* genes than automated MQ bins, and therefore genomic data was recovered for only about 70% of the communities after correction and polishing. Together with the fact that prior SR sequencing and assembly of the NLR yielded 49 unique *rpoB* genes[79], this demonstrates the relative weakness of LR assemblies ability to capture the same breadth and depth of community members, presumably due to a much more shallow sequencing depth.

As expected based on the technological differences between LR and SR sequencing platforms, mean depths of *rpoB*-containing contigs were slightly higher for SRs than LRs, with 27.9 and 21.5 in the OLR, and 23.5 and 19.5 in the NLR, respectively (Fig. S5). Furthermore, both LR and SR recruitment depths were lower for reads mapped to the opposite bioreactor's assembly, indicating that while there is some overlap as expected given that both reactor systems were inoculated with biomass from the same source, the communities were distinct. In fact, there was a strong correlation between the LR and SR depths themselves (adjusted $R^2$ >0.97, p-value <<0.05, data not shown). However, the mean depth of both LRs and SRs of the same bioreactor for *rpoB*-containing contigs was reduced by SR polishing by 12.5% and 15.2%

respectively. This phenomenon possibly occurred because lower abundance community members with more erroneously assembled sequences required SR polishing for their *rpoB* genes to be identified. However, we also note the possibility that reads were incorrectly aligned to erroneous sequences causing depth variation, but this was not explored further. Additionally, while SR depth was greater than LR for the same bioreactor assembly, more LRs than SRs from the other bioreactor could be recruited to the opposite bioreactor's assembly. This, like the ability to recruit reads from the opposite bioreactor generally, may be caused by the presence of similar but distinct strains in the two reactor systems and more relaxed alignment thresholds for LR compared to SR recruitment.

Both LR correction and SR polishing caused substantial shifts in the observed bioreactor communities. On the one hand, LR correction was noisy and communities remained separated rather than forming tight clusters (Fig. 5). On the other hand, SR polishing without and with LR correction led to major community shifts followed by convergence. There was a dependence on both bioreactor and assembler, for example Canu assemblies appear to overlap more after some LR correction than without, which was not as apparent for the Flye assemblies. This effect possibly was caused by the lower diversity captured by Canu assemblers rather than a greater improvement by LR correction of Canu than Flye assemblies. Furthermore, the 10 iterations of LR correction for the OLR Flye assembly deviated substantially from the rest of the assemblies but still converged, albeit on its own path, during SR polishing. Thus, while LR correction does affect the recovery efficiency of the community members, SR polishing again causes the greatest and least noisy changes that yield the most consistent results and likely best representation of the microbial community present.

*Additional information on automated bin recovery*

Automated binning was dependent on both bioreactor and assembler, with minimal improvement from LR correction and major improvements from SR polishing. Initial LR assemblies led to the recovery of between 2 and 16 medium-quality or better (MQ) bacterial genomic bins. Both bioreactor and assembler had a major influence, with approximately 1.9-fold more MQ bins recovered from the initial NLR assemblies compared to the OLR assemblies, and 2.5-fold more MQ bins recovered from Flye compared to Canu assemblies. On average, over all LR correction iterations, the MQ bin yield increased by less than 3 to 12.3 (Fig. S6). Consistent with other results, LR correction remained noisy after the first iteration: on average one iteration of LR correction allowed 3 additional MQ bins to be recovered, increased the number of contigs in MQ bins from 33.8 to 75.8, and increased the bps in MQ bins from 39,179,008 to 51,993,034. These improvements decreased after the second iteration, reached similar values again after the third iteration as after the first, and were overall somewhat unstable during LR correction (Fig. 4, Fig. S6). SR polishing without and with LR correction on average allowed 17.6 and 19.1 MQ automated bins to be recovered, yielded 188 and 297 contigs comprising 72,619,837 and 78,451,859 bps in MQ bins, and lead to an increase in completeness scores to 82.9% and 79.7%, respectively. Again, the majority of improvement occurred within the first 3 iterations of SR polishing, with diminishing returns after subsequent iterations (Fig. 4, Fig. S6). We acknowledge that the mean completeness scores of MQ bins were lower from assemblies that were both LR corrected and SR polished rather than SR polished alone. Given the increase in MQ bin count, as well as the number of contigs and bps in MQ bins, it is most likely that lower quality bins or inaccurate contigs improved to meet the MQ bin threshold, thereby still achieving a better representation of the microbial community.

Reference-agnostic bacterial marker gene redundancy at the assembly levels estimated using CheckM followed similar patterns as *rpoB* gene recovery. First, initial LR assemblies tended to have the lowest redundancy, *i.e.*, the fewest number of the same marker gene in the assembly (Fig. S6). Furthermore, the Canu assembly of the OLR lacked 5/104 marker genes altogether, while at least two copies of each of the 104 marker genes were identified in the Flye assembly of the NLR. Second, LR correction alone was noisy and insufficient to increase redundancy to 5 or more copies of each gene; for all assemblies except the NLR Flye assembly, the average number of genes present in 5+ copies was 99.2/104 after LR correction compared to 97/104 prior to LR correction. Third, SR polishing led to the identification of at least five copies of each marker gene and increased redundancy to essentially the maximum recorded by the third iteration, regardless of prior LR correction (Fig. S6). Fourth, preceding SR polishing by any LR correction increased the total estimated redundancy, *i.e.*, CheckM contamination score, compared to SR polishing of the initial assemblies from 2,558% to 2,670% on average, demonstrating a benefit of at least some LR correction to increase marker gene redundancy.

Intriguingly, we noted a strong correlation (adjusted $R^2 \geqq 0.85$, p-value <<0.05) between the observed redundancy of the full assembly, *i.e.*, the sum of CheckM completion and contamination scores, and the number of medium-quality or better automated bins. This suggests that assembly redundancy may be used as a predictor for genome recovery (Fig. S7). We do acknowledge that this relationship was largely driven by large improvements as a result of SR polishing, as indicated by the data typically clustering into two distinct groups based on the iterations of LR correction. Statistical testing with fewer SR polishing iterations or different assembly characteristics yielded sometimes weaker (adjusted $R^2 \geq 0.7$) but still significant (p-value <0.05) correlations (data not shown), indicating an overall robust relationship. Taken together, both marker gene redundancy for the entire assembly and automated bin quality vastly improved by SR polishing with some benefit also gained from LR correction.

*Additional information on gene fragmentation*

Initial LR assemblies contained, on average, 329,650 coding genes and 136,613,267 bps in these coding genes, or approximately 76% of all assembled bps. There were clear differences in the coding gene contents of the assemblies due to both bioreactor and assembler. The OLR assemblies contained on average 281,935 genes with 114,083,729 bps in them while the NLR assemblies contained 377,364 genes with 159,142,804 bps. Canu assemblies contained 313,901 genes with 122,320,542 bps in them while the Flye assembles contained 345,398 with 150,905,992 bps. These differences are consistent with the assembly contiguity and the greater gene yields in the NLR assemblies over the OLR assemblies are likely due to the greater sequencing depth, while the higher yields from Flye over Canu assemblies suggest that Flye was able to reconstruct higher-quality assemblies.

Gene fragmentation was slightly reduced by LR correction and drastically by SR polishing. LR correction alone led to only moderate improvements in gene fragmentation, on average reducing the gene count to 325,330 and increasing bps in genes to 139,661,940, or approximately a 1.3% and 2.2% change respectively. On the other hand, SR polishing alone on average reduced the gene count to 266,977 and simultaneously increased the bps in genes to 153,102,694, or approximately 19% and 12% change, respectively, comprising approximately 85% of all assembled bps on average. This improvement was slightly more pronounced when implementing SR polishing after any LR correction, reducing the gene count to 265,953 and increasing the bps in genes to 154,819,887. Across several fractions of the assemblies, *i.e.*, circular or long contigs that may represent essentially complete bacterial replicons (chromosomes or plasmids), and MQ bins, the median quantity of coding genes and bps in coding genes normalized to the Mbps contig length quickly saturates after about two to four SR polishing iterations but never due to LR correction alone (Fig. S8). Therefore, not only does the improvement occur throughout the assembly, but these improvements are also apparent, if not to a greater magnitude, for subsets of the assemblies most tractable for downstream analyses.

Additionally, these same trends hold true when looking at the gene density of the assemblies. After a single iteration, the median total number of genes per contig normalized to Mbp of sequence increased on average by 38 (~2.5%) genes due to LR correction but decreased on average by 98 (~6.3%) genes due to SR polishing (Fig. S9). Again, after one iteration, the median total bps in genes per contig normalized to Mbp of sequence increased due to both LR correction and SR polishing, by 14,477 (~2.3%) and 22,722 (~3.5%) bps on average, respectively. Over all iterations of error fixing, LR correction alone led to an average increase of 33 genes and 14,184 bps in genes, while SR polishing without LR correction led to the largest decrease in genes (166, ~10.7%) and a moderate increase in bps in genes (31,486, ~4.9%) and SR polishing with at least some LR correction led to a large decrease in genes (145, ~9.4%) and the largest increase in bps in genes (49,332, ~7.7%).

According to BUSO, which identifies 124 bacterial marker genes and assesses their fragmentation and redundancy, all LR assemblies initially contained fragmented bacterial marker genes, which were not resolved by LR correction alone. Consistent with general assembly statistics, initial Flye assemblies contained on average 3 additional complete (*i.e.*, not fragmented) and 22 additional recovered (*i.e.*, not missing) marker genes than the initial Canu assemblies (Fig. S8). Similarly, there were on average 1.5 more complete and 24 more recovered marker genes in the NLR compared to the OLR, again likely due to the greater read depth. Even one iteration of LR correction increased complete and recovered bacterial marker genes, by an average of 2 and 10, respectively, but it was also evident that up to 10 iterations of LR correction could not fully resolve these errors and led to noisy patterns. These observations were at least partially expected because LR correction still uses the high error rate LRs and is typically used to fix slightly larger scale, *i.e.*, structural, errors.

On the other hand, SR polishing effectively resolved gene fragmentation when viewed as an aggregate over the entire assembly. Without any LR correction, only one iteration of SR polishing was sufficient to recover at least one complete copy of all tracked marker genes. Furthermore, this trend was consistent after every LR correction step tested, suggesting that the SR polishing outcomes are more stable than LR correction, which is expected because they tend to fix smaller scale, *i.e.*, single or few bps errors. We additionally note that for the Flye assembly of the OLR, more markers were complete and duplicated with 0 LR correction steps than with 2, 5, or 10 iterations, in contrast to the other three LR assemblies. This may be the result of the LR corrections resolving an inaccuracy, in which a large redundant sequence was collapsed (more evidence below), but this phenomenon was not extensively followed. In summary, it was clear that only a few iterations of SR polishing were sufficient to lead to the reduction of fragmented marker genes, but LR correction demonstrated diminished returns and noisy patterns.

Corroborating the trends observed in BUSCO, the broader reference comparison with IDEEL showed that initial assemblies contained many fragmented genes, but SR polishing led to major improvements in gene length. On average, only about 6.8% of coding genes in the initial assemblies were within 5% of the length of their most similar reference sequence (Fig. S10). LR correction alone increased this proportion to only 7.9% of coding genes, but just two rounds of SR polishing increased the proportion of coding genes with lengths within 5% of their most similar reference to 24.3%. There was a slight improvement by the third iteration of SR polishing to 24.8% and with subsequent SR polishing, diminishing returns to a maximum of 25.0%. Corroborating the shift towards reference length, moderately fragmented (≤50% reference length) and highly fragmented (≤10%) genes in the assemblies decreased from 55.5% to 37.6% and from 5.8% to 3.7% respectively.

*Additional information on short read recruitment*

The SR recruitment of initial LR assemblies recapitulated many of the other observed patterns, for example major differences between assemblers but less pronounced differences between bioreactors. The OLR assemblies recruited 451,908,712 SR bps; 4.88% of SRs were ambiguously aligned, and 1,378 contigs recruited no SRs. The NLR assemblies recruited 492,299,062 SR bps, with 3.38% of SRs ambiguously aligned, and 2,180 contigs not recruiting any SRs. Flye assemblies recruited 617,179,142 SR bps, with 4.76% of SRs ambiguously aligned, and 405 contigs not recruiting any SRs, while the Canu assemblies recruited 327,028,632 SR bps, with 3.50% of SRs ambiguously aligned, and 3,154 contigs not recruiting any SRs. Thus, sequencing depth had a smaller effect on SR recruitment than assembler. While Flye assemblies were able to recruit more SR and to more of the contigs, there were also more ambiguously aligned reads, suggesting that much of the assembly may be higher quality and contain similar yet distinct strains, or could contain a large duplication.

Consistent with prior results and expectations, LR correction tended to make only minor improvements to SR recruitment, but SR polishing led to major improvements. After only LR correction of the Flye assemblies, the total SR bps recruited changed negligibly, *i.e.*, <5% increase or decrease, but increased to 596,399,632 bps for Canu assemblies on average. After SR polishing, Canu assemblies on average recruited 1,430,808,700 total SR bps and Flye assemblies 1,456,688,314 total SR bps, representing a 4.38-fold and 2.36-fold increase, respectively.

Normalizing for assembly sizes may increase resolution to help assess the distribution of SR recruitment; the median SR bps recruited normalized to contig length per Mbp for Flye assemblies increased from 29,717 to 37,170 due to LR correction alone, which is a much larger apparent difference (25.1%) than total SR bps recruited. However, the median SR bps recruited normalized to contig length per Mbp for Canu assemblies did not increase from 0 due to LR correction, highlighting issues with both Canu assemblies and this analytical approach. Fortunately, LR correction led to 31,453 greater median SR recruited bps per contig per Mbp after subsequent SR polishing (93,579) compared to SR polishing alone (62,126), indicating both a benefit of some LR correction prior to SR polishing and that SR polishing was the major driver of improved SR recruitment. Similarly, for Flye assemblies, the median SR bps per contig per Mbp increased to 223,421 with SR polishing without LR correction and to 249,896 with LR correction preceding SR polishing, or approximately 7.52- and 8.41-fold increases, respectively. For all assemblies, SR recruitment largely saturated after the third iteration of SR polishing, leading to an average 3.06-fold increase from 472,103,888 to 1,445,120,660 total SR bps recruited, and 10.6-fold increase from 14,859 to 157,255 median SR bps per contig per Mbp for all assemblies.

The ambiguity of SR recruitment also followed similar patterns as the total bps in reads per contig normalized to Mbp. Assemblies typically started with 3.2-3.6% of the total reads mapping ambiguously, but 6.3% in the Flye assembly of the OLR, presumably due to a large-scale duplication that was collapsed by minimal LR correction or reconstruction of multiple similar strains. LR correction and SR polishing alone both reduced the fraction of ambiguously mapped reads by approximately 30-50%, but only when combining both LR correction and SR polishing was ambiguity reduced maximally by about 70%, or to 1.2-1.9% of total mapped reads. Most of this ambiguity reduction occurred within the first one or two rounds of SR correction. Therefore, given both the overall increase in SR recruitment and simultaneous decrease in SR ambiguity, at least one iteration of LR correction and at least two iterations of SR polishing vastly improved the accuracy of the assembly. However, it should be noted that we observed diminishing returns, especially beyond 5 iterations of SR polishing.

The number of contigs without any SR recruitment generally decreased in all correction approaches tested, but again primarily due to SR polishing. Canu assemblies, initially containing more contigs than Flye assemblies (see prior section), also contained more contigs that did not recruit any SRs (Canu assemblies: 2,408/4,187 [OLR] and 3,899/6,951 [NLR]; Flye assemblies: 348/1,487 [OLR] and 462/2,301 [NLR]), with fractions of contigs not recruiting any SR of 57% and 22% for the Canu and Flye assemblies, respectively. On average over all ten iterations of LR correction for both bioreactors and assemblers, the number of contigs that did not recruit any SR decreased from 1,779 to 1,630 (8% reduction). Without LR correction, SR polishing reduced these to 1,268 (29% reduction), and to 1,184 (33% reduction) with at least two preceding LR correction iterations. These data demonstrated that SR polishing causes large decreases in the number of contigs recruiting no SRs and a slight benefit of some LR correction prior to SR polishing. The largest improvements occurred after just one iteration of SR polishing, with only marginal improvements beyond four iterations regardless of bioreactor or assembler.

(14) Kolmogorov, M.; Bickhart, D. M.; Behsaz, B.; Gurevich, A.; Rayko, M.; Shin, S. B.; Kuhn, K.; Yuan, J.; Polevikov, E.; Smith, T. P. L.; Pevzner, P. A. metaFlye: Scalable Long-Read Metagenome Assembly Using Repeat Graphs.

*Nat Methods* **2020**, *17* (11), 1103–1110. https://doi.org/10.1038/s41592-020-00971-x.

(26) Van Goethem, M. W.; Osborn, A. R.; Bowen, B. P.; Andeer, P. F.; Swenson, T. L.; Clum, A.; Riley, R.; He, G.; Koriabine, M.; Sandor, L.; Yan, M.; Daum, C. G.; Yoshinaga, Y.; Makhalanyane, T. P.; Garcia-Pichel, F.; Visel, A.;

Pennacchio, L. A.; O'Malley, R. C.; Northen, T. R. Long-Read Metagenomics of Soil Communities Reveals Phylum-Specific Secondary Metabolite Dynamics. *Commun Biol* **2021**, *4* (1), 1–10. https://doi.org/10.1038/s42003-021-02809-4.

(39) Warren, R. L.; Coombe, L.; Mohamadi, H.; Zhang, J.; Jaquish, B.; Isabel, N.; Jones, S. J. M.; Bousquet, J.; Bohlmann, J.; Birol, I. ntEdit: Scalable Genome Sequence Polishing. *Bioinformatics* **2019**, *35* (21), 4430–4432. https://doi.org/10.1093/bioinformatics/btz400.
(40) Shafin, K.; Pesout, T.; Lorig-Roach, R.; Haukness, M.; Olsen, H. E.; Bosworth, C.; Armstrong, J.; Tigyi, K.; Maurer, N.; Koren, S.; Sedlazeck, F. J.; Marschall, T.; Mayes, S.; Costa, V.; Zook, J. M.; Liu, K. J.; Kilburn, D.; Sorensen, M.; Munson, K. M.; Vollger, M. R.; Monlong, J.; Garrison, E.; Eichler, E. E.; Salama, S.; Haussler, D.; Green, R. E.; Akeson, M.; Phillippy, A.; Miga, K. H.; Carnevali, P.; Jain, M.; Paten, B. Nanopore Sequencing and the Shasta Toolkit Enable Efficient de Novo Assembly of Eleven Human Genomes. *Nat Biotechnol* **2020**, *38* (9), 1044–1053. https://doi.org/10.1038/s41587-020-0503-6.
(41) Li, H. Minimap and Miniasm: Fast Mapping and de Novo Assembly for Noisy Long Sequences. *Bioinformatics* **2016**, *32* (14), 2103–2110. https://doi.org/10.1093/bioinformatics/btw152.
(42) Hu, J.; Wang, Z.; Sun, Z.; Hu, B.; Ayoola, A. O.; Liang, F.; Li, J.; Sandoval, J. R.; Cooper, D. N.; Ye, K.; Ruan, J.; Xiao, C.-L.; Wang, D.-P.; Wu, D.-D.; Wang, S. An Efficient Error Correction and Accurate Assembly Tool for Noisy Long Reads. bioRxiv March 12, 2023, p 2023.03.09.531669. https://doi.org/10.1101/2023.03.09.531669.
(43) Vaser, R.; Šikić, M. Time- and Memory-Efficient Genome Assembly with Raven. *Nat Comput Sci* **2021**, *1* (5), 332–336. https://doi.org/10.1038/s43588-021-00073-4.
(44) Lv, X.; Chen, Z.; Lu, Y.; Yang, Y. An End-to-End Oxford Nanopore Basecaller Using Convolution-Augmented Transformer. bioRxiv November 10, 2020, p 2020.11.09.374165. https://doi.org/10.1101/2020.11.09.374165.
(45) Miculinić, N.; Ratković, M.; Šikić, M. MinCall - MinION End2end Convolutional Deep Learning Basecaller. arXiv April 22, 2019. https://doi.org/10.48550/arXiv.1904.10337.
(46) Zeng, J.; Cai, H.; Peng, H.; Wang, H.; Zhang, Y.; Akutsu, T. Causalcall: Nanopore Basecalling Using a Temporal Convolutional Network. *Frontiers in Genetics* **2020**, *10*.
(47) Huang, N.; Nie, F.; Ni, P.; Luo, F.; Wang, J. SACall: A Neural Network Basecaller for Oxford Nanopore Sequencing Data Based on Self-Attention Mechanism. *IEEE/ACM Transactions on Computational Biology and Bioinformatics* **2022**, *19* (1), 614–623. https://doi.org/10.1109/TCBB.2020.3039244.
(48) Konishi, H.; Yamaguchi, R.; Yamaguchi, K.; Furukawa, Y.; Imoto, S. Halcyon: An Accurate Basecaller Exploiting an Encoder–Decoder Model with Monotonic Attention. *Bioinformatics* **2021**, *37* (9), 1211–1217. https://doi.org/10.1093/bioinformatics/btaa953.
(49) Wu, Y.-W.; Tang, Y.-H.; Tringe, S. G.; Simmons, B. A.; Singer, S. W. MaxBin: An Automated Binning Method to Recover Individual Genomes from Metagenomes Using an Expectation-Maximization Algorithm. *Microbiome* **2014**, *2* (1), 26. https://doi.org/10.1186/2049-2618-2-26.
(50) Uritskiy, G. V.; DiRuggiero, J.; Taylor, J. MetaWRAP—a Flexible Pipeline for Genome-Resolved Metagenomic Data Analysis. *Microbiome* **2018**, *6* (1), 158. https://doi.org/10.1186/s40168-018-0541-1.
(51) Wang, Z.; Huang, P.; You, R.; Sun, F.; Zhu, S. MetaBinner: A High-Performance and Stand-Alone Ensemble Binning Method to Recover Individual

Genomes from Complex Microbial Communities. *Genome Biology* **2023**, *24* (1), 1. https://doi.org/10.1186/s13059-022-02832-6.
(52) *UniteM/unitem_ms.draft.pdf at master · donovan-h-parks/UniteM*. GitHub. https://github.com/donovan-h-parks/UniteM/blob/master/unitem_ms.draft.pdf (accessed 2023-12-13).
(53) Imelfort, M.; Parks, D.; Woodcroft, B. J.; Dennis, P.; Hugenholtz, P.; Tyson, G. W. GroopM: An Automated Tool for the Recovery of Population Genomes from Related Metagenomes. *PeerJ* **2014**, *2*, e603. https://doi.org/10.7717/peerj.603.
(54) Donato, L.; Scimone, C.; Rinaldi, C.; D'Angelo, R.; Sidoti, A. New Evaluation Methods of Read Mapping by 17 Aligners on Simulated and Empirical NGS Data: An Updated Comparison of DNA- and RNA-Seq Data from Illumina and Ion Torrent Technologies. *Neural Comput & Applic* **2021**, *33* (22), 15669–15692. https://doi.org/10.1007/s00521-021-06188-z.
(55) Otto, C.; Stadler, P. F.; Hoffmann, S. Lacking Alignments? The next-Generation Sequencing Mapper Segemehl Revisited. *Bioinformatics* **2014**, *30* (13), 1837–1843. https://doi.org/10.1093/bioinformatics/btu146.
(56) Dadi, T. H.; Siragusa, E.; Piro, V. C.; Andrusch, A.; Seiler, E.; Renard, B. Y.; Reinert, K. DREAM-Yara: An Exact Read Mapper for Very Large Databases with Short Update Time. *Bioinformatics* **2018**, *34* (17), i766–i772. https://doi.org/10.1093/bioinformatics/bty567.
(57) Kim, D.; Paggi, J. M.; Park, C.; Bennett, C.; Salzberg, S. L. Graph-Based Genome Alignment and Genotyping with HISAT2 and HISAT-Genotype. *Nat Biotechnol* **2019**, *37* (8), 907–915. https://doi.org/10.1038/s41587-019-0201-4.
(58) Li, H.; Durbin, R. Fast and Accurate Short Read Alignment with Burrows–Wheeler Transform. *Bioinformatics* **2009**, *25* (14), 1754–1760. https://doi.org/10.1093/bioinformatics/btp324.
(59) Langmead, B.; Salzberg, S. L. Fast Gapped-Read Alignment with Bowtie 2. *Nat Methods* **2012**, *9* (4), 357–359. https://doi.org/10.1038/nmeth.1923.
(60) Marco-Sola, S.; Sammeth, M.; Guigó, R.; Ribeca, P. The GEM Mapper: Fast, Accurate and Versatile Alignment by Filtration. *Nat Methods* **2012**, *9* (12), 1185–1188. https://doi.org/10.1038/nmeth.2221.
(61) LoTempio, J.; Délot, E.; Vilain, E. Benchmarking Long-Read Genome Sequence Alignment Tools for Human Genomics Applications. bioRxiv June 20, 2023, p 2021.07.09.451840. https://doi.org/10.1101/2021.07.09.451840.
(62) Yildiz, G.; Zanini, S. F.; Afsharyan, N. P.; Obermeier, C.; Snowdon, R. J.; Golicz, A. A. Benchmarking Oxford Nanopore Read Alignment-Based Insertion and Deletion Detection in Crop Plant Genomes. *The Plant Genome* **2023**, *16* (2), e20314. https://doi.org/10.1002/tpg2.20314.
(63) Marić, J.; Sović, I.; Križanović, K.; Nagarajan, N.; Šikić, M. Graphmap2 - Splice-Aware RNA-Seq Mapper for Long Reads. bioRxiv July 30, 2019, p 720458. https://doi.org/10.1101/720458.
(64) Ren, J.; Chaisson, M. J. P. Lra: A Long Read Aligner for Sequences and Contigs. *PLOS Computational Biology* **2021**, *17* (6), e1009078. https://doi.org/10.1371/journal.pcbi.1009078.
(65) Sedlazeck, F. J.; Rescheneder, P.; Smolka, M.; Fang, H.; Nattestad, M.; von Haeseler, A.; Schatz, M. C. Accurate Detection of Complex Structural Variations Using Single-Molecule Sequencing. *Nat Methods* **2018**, *15* (6), 461–468. https://doi.org/10.1038/s41592-018-0001-7.

(102) Oksanen, J.; Simpson, G. L.; Blanchet, F. G.; Kindt, R.; Legendre, P.; Minchin, P. R.; O'Hara, R. B.; Solymos, P.; Stevens, M. H. H.; Szoecs, E.; Wagner, H.; Barbour, M.; Bedward, M.; Bolker, B.; Borcard, D.; Carvalho, G.; Chirico, M.; Caceres, M. D.; Durand, S.; Evangelista, H. B. A.; FitzJohn, R.; Friendly, M.; Furneaux, B.; Hannigan, G.; Hill, M. O.; Lahti, L.; McGlinn, D.; Ouellette, M.-H.; Cunha, E. R.; Smith, T.; Stier, A.; Braak, C. J. F. T.; Weedon, J. *Vegan: Community Ecology Package*; 2022.

(103) R Core Team. *R: A Language and Environment for Statistical Computing*; R Foundation for Statistical Computing: Vienna, Austria, 2021.
