## Supplementary material for "Simple, reference-independent analyses help optimize hybrid assembly of microbial community metagenomes": Figure S1

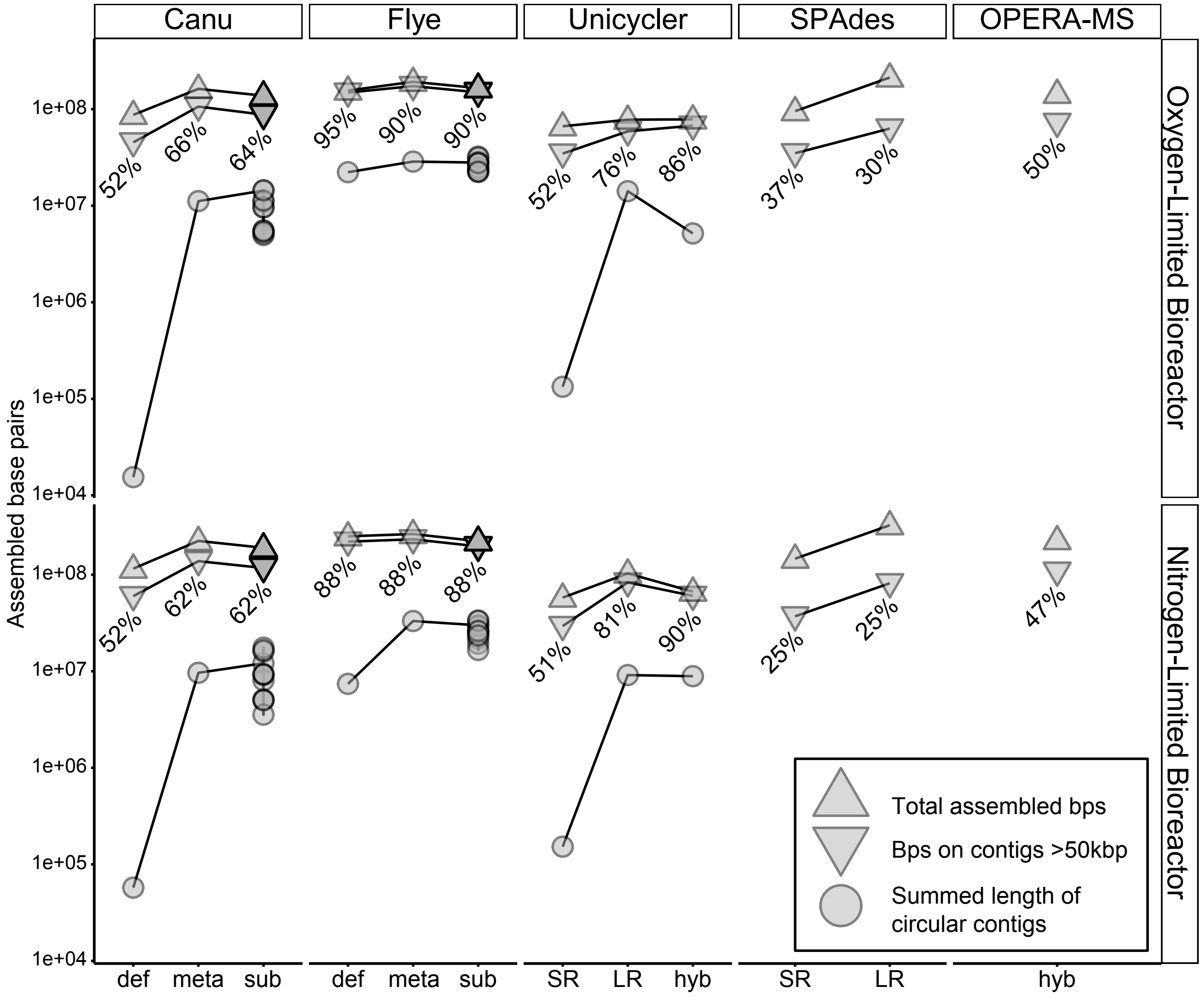


**Figure S1. Plot of assembly yields from different programs for each bioreactor.**
In each panel, separated vertically by reactor and horizontally by assembler, the total assembled bps (upward-pointing triangles), bps on contigs >50 Kbp (downward pointing triangles), and bps in circular contigs (circles) are shown on the y-axis and grouped by the assembly type or setting on the x-axis. Numbers indicate the percentage of total bps on the larger (>50 Kbps) contigs. Abbreviations are as follows: def, default settings; meta, metagenome-optimized settings; sub, assembly of sub-sampled reads; SR, short read only metagenomic assembly; LR, long read only assembly; hyb, hybrid assembly.
