## Supplementary material for "Simple, reference-independent analyses help optimize hybrid assembly of microbial community metagenomes": Figure S2

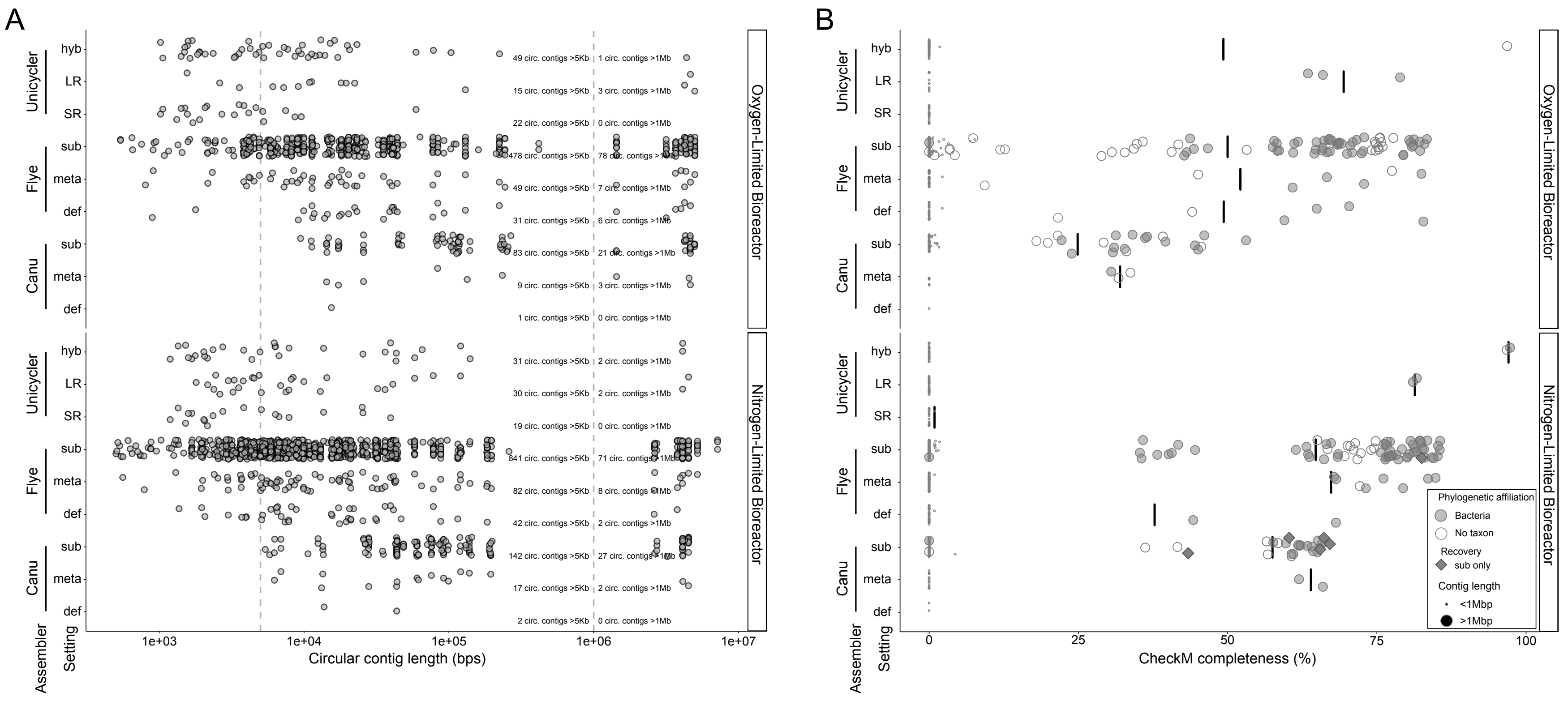


**Figure S2. Jittered plots of circular contig recovery from different assembly programs for both bioreactors.** The two bioreactors are separated over the vertical panels. (A) Length distribution of circular contigs for each assembler. Vertical dashed lines indicate common thresholds for considering complete genomes or chromosomes, >5 Kbp for plasmids and phages, and >1 Mbp for microbial genomes, though exceptions exist. The number of contigs passing these thresholds are indicated next to the right-most vertical line. (B) Completion of circular contigs estimated using lineage-specific phylogenetic markers. Filled circles indicate a circular contig with bacterial phylogenetic markers, empty circles indicate a circular contig lacking sufficient data to assign to the bacterial domain, and filled diamonds indicate a bacterial lineage only circularized in sub-sampled assemblies. Black horizontal bar represents the mean of circular contigs with non-zero completion estimates. Abbreviations are as follows: def, default settings; meta, metagenome-optimized settings; sub, assembly of sub-sampled reads; SR, short read only metagenomic assembly; LR, long read only assembly; hyb, hybrid assembly.
