## Supplementary material for "Simple, reference-independent analyses help optimize hybrid assembly of microbial community metagenomes": Figure S3

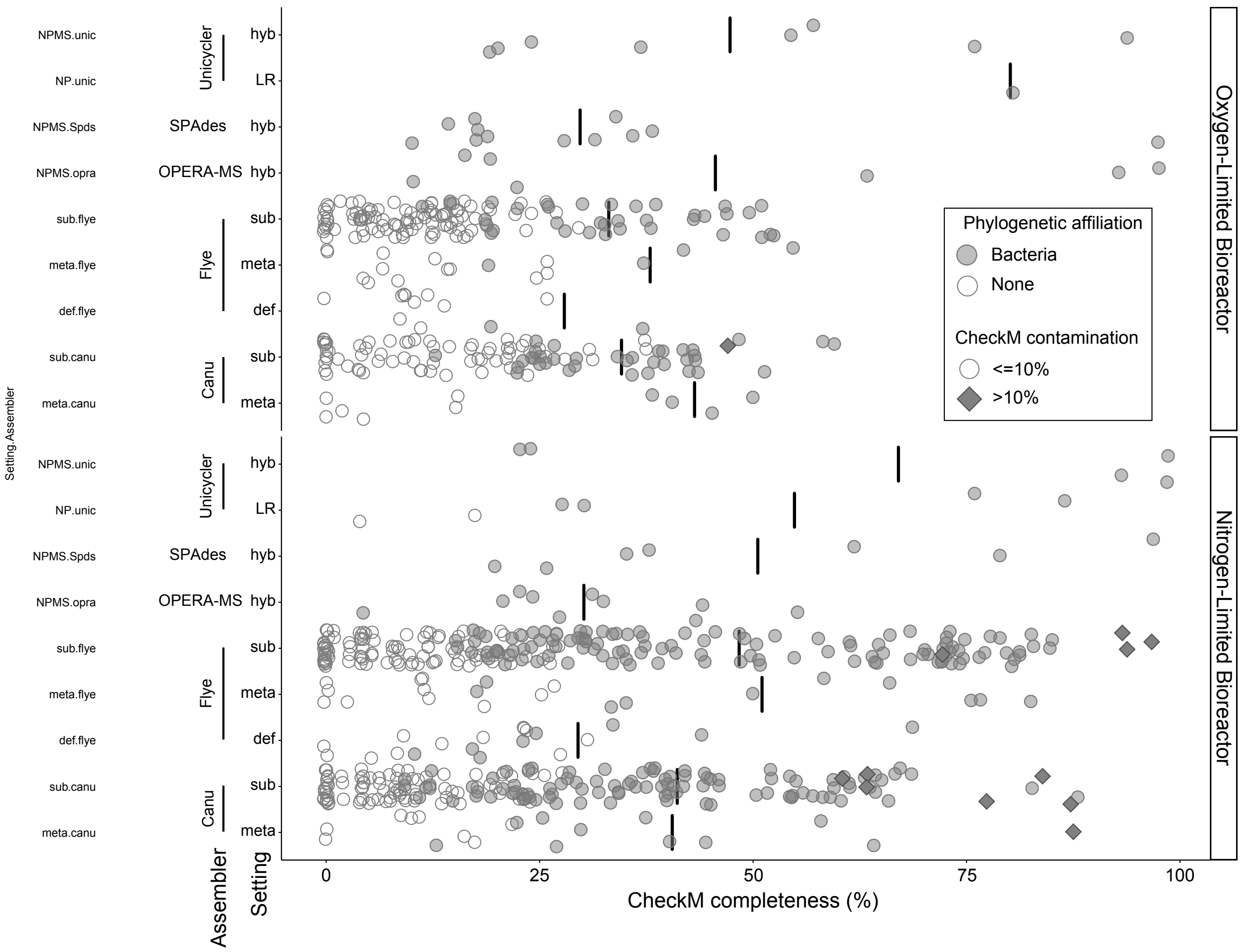


**Figure S3. Jittered plot of non-circular, large (>1 Mbp) contig recovery from different assembly programs for both bioreactors.**
The two bioreactors are separated over the vertical panels. Filled circles indicate >1 Mbp contigs containing bacterial phylogenetic markers, empty circles indicate >1 Mbp contigs lacking sufficient data to assign to the bacterial domain, and filled diamonds indicate a contig with >10% redundancy of bacterial phylogenetic markers. Black horizontal bars represent the mean of >1 Mbp contigs with sufficient data to be affiliated with the bacterial domain. Abbreviations are as follows: def, default settings; meta, metagenome-optimized settings; sub, assembly of sub-sampled reads; SR, short read only metagenomic assembly; LR, long read only assembly; hyb, hybrid assembly.
