## Supplementary material for "Simple, reference-independent analyses help optimize hybrid assembly of microbial community metagenomes": Figure S4

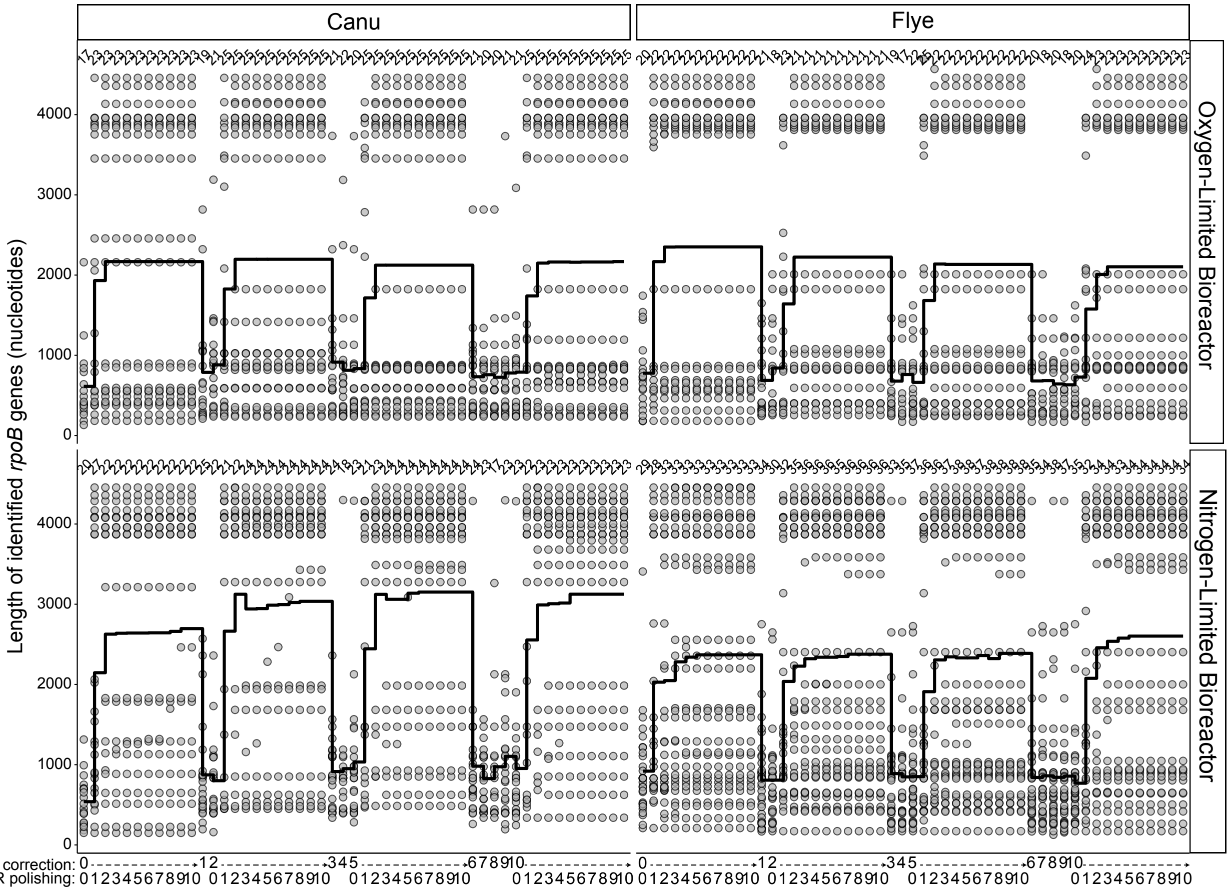


**Figure S4. Stair plot of assembled RNA polymerase subunit B (*rpoB*) gene lengths for each bioreactor and Long Read (LR) assembly throughout the iterative LR correction and Short Read (SR) polishing processes.**

The two bioreactors are separated over vertical panels, the two LR assemblers over the horizontal panels. The LR correction and SR polishing iterations are spread across the x-axis so that the ten SR polishing steps are immediately to the right of the preceding LR correction step. Points are the length of each *rpoB* gene in the assembly, with the black line showing the mean length for the assembly and the text above the points indicating the count.
