## Supplementary material for "Simple, reference-independent analyses help optimize hybrid assembly of microbial community metagenomes": Figure S5

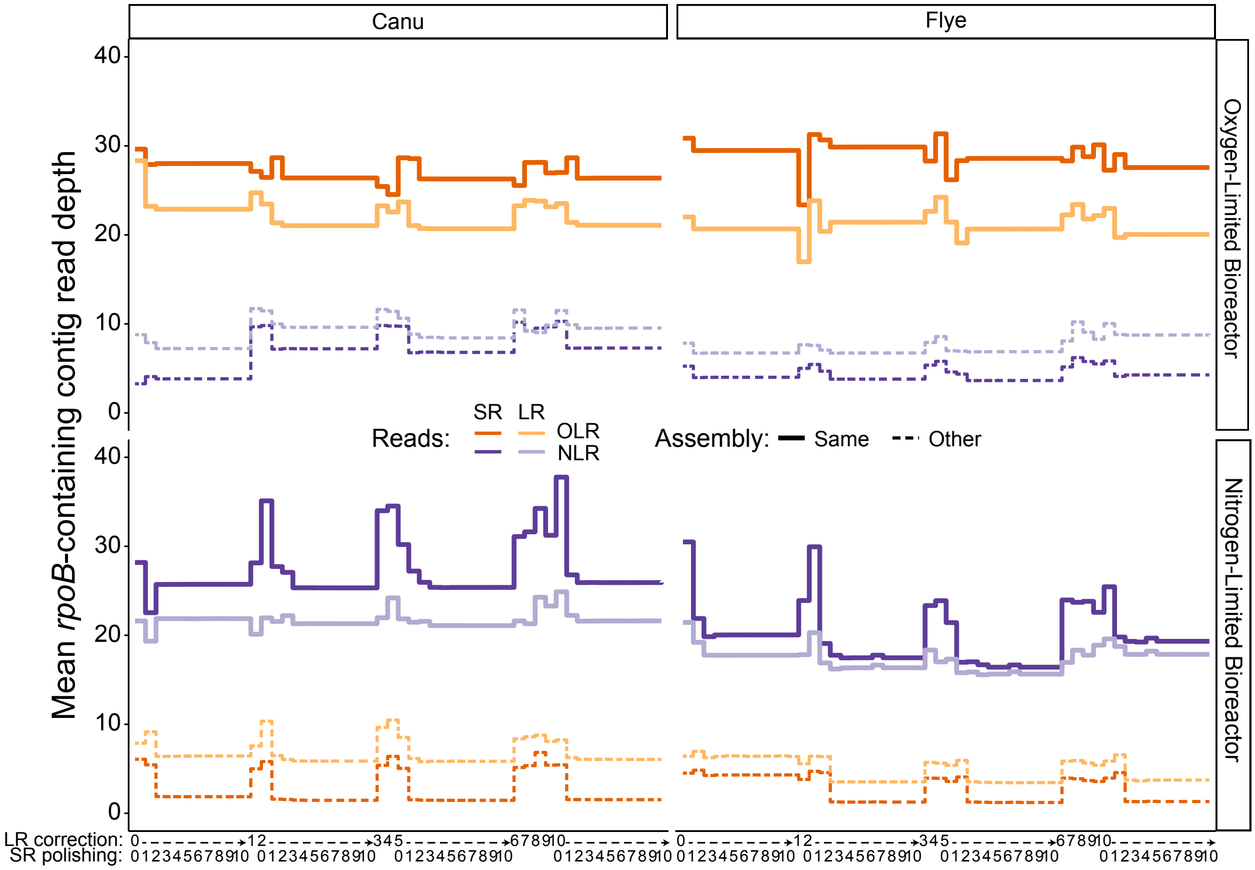


**Figure S5. Stair plot of assembled RNA polymerase subunit B (*rpoB*) contig read depths for each bioreactor and Long Read (LR) assembly throughout the iterative LR correction and Short Read (SR) polishing processes.**
The two bioreactors are separated over vertical panels, the two LR assemblers over the horizontal panels. The LR correction and SR polishing iterations are spread across the x-axis so that the ten SR polishing steps are immediately to the right of the preceding LR correction step. Colored lines indicate the read source used for mean coverage calculation.
