## Supplementary material for "Simple, reference-independent analyses help optimize hybrid assembly of microbial community metagenomes": Figure S6

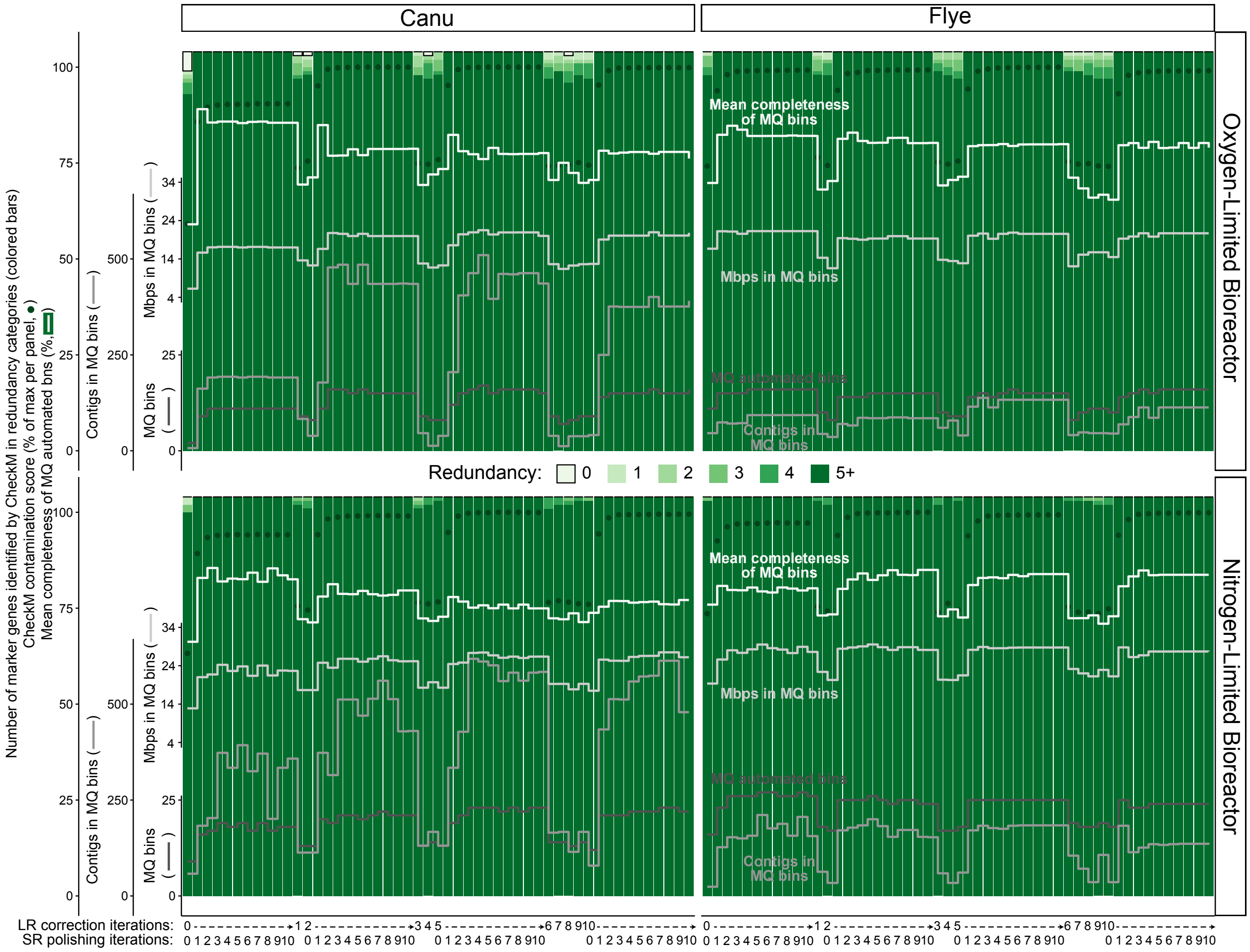


**Figure S6. Stacked bar plot of CheckM marker gene redundancy overlaid with stair plots of medium-quality or better (MQ) automated bin information for each bioreactor and Long Read (LR) assembly throughout the iterative LR correction and Short Read (SR) polishing processes.**

The two bioreactors are separated over vertical panels, the two LR assemblers over the horizontal panels. The long read (LR) correction and SR polishing iterations are spread across the x-axis so that the ten SR polishing steps are immediately to the right of the preceding LR correction step. Colored bars show the CheckM copy number estimates (maximum reported is “5+”) for the entire assembly, the gray lines indicate additional information for automated bins scaled to overlay the bars for several fractions of the assemblies throughout the LR correction and SR polishing iterations.
