## Supplementary material for "Simple, reference-independent analyses help optimize hybrid assembly of microbial community metagenomes": Figure S11

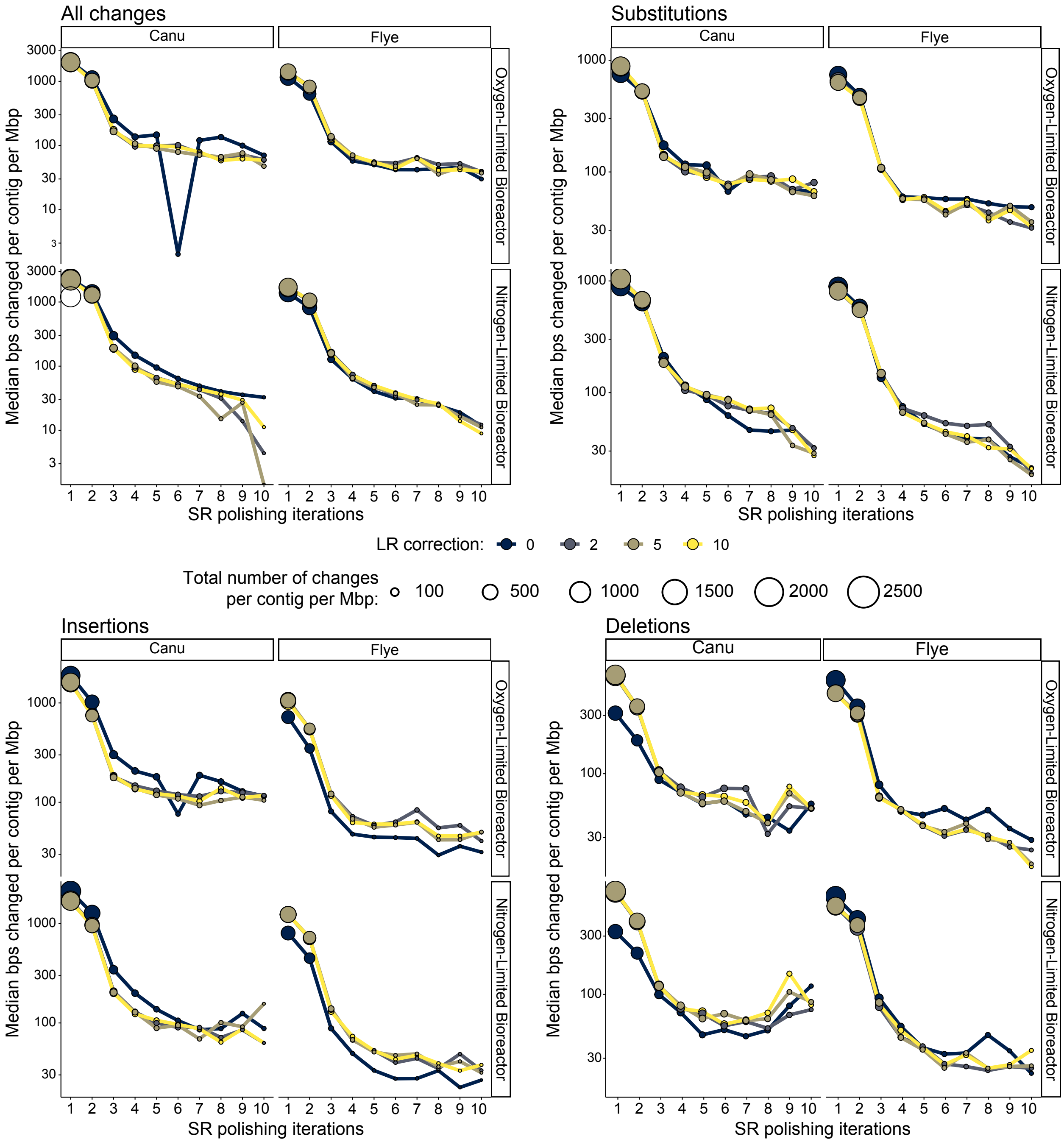


**Figure S11. Curves of extent of changes made during Short Read (SR) polishing for each bioreactor and Long Read (LR) assembly throughout the iterative LR correction and SR polishing processes.**

(A) All changes made during SR polishing. (B) Substitutions made during SR polishing. (C) Insertions made during SR polishing. (D) Deletions made during SR polishing. Within each main panel the two bioreactors are separated over vertical sub-panels, and the two LR assemblers are separated over the horizontal sub-panels. The x-axis shows the SR polishing iterations. Points are the median changes per contig per Mbp sequence for each assembly and are area-scaled by the median total bps changed per contig per Mbp sequence. Each point is colored by the SR polishing iteration, with colored lines connecting the points indicating the LR correction iteration.
