## Supplementary material for "Simple, reference-independent analyses help optimize hybrid assembly of microbial community metagenomes": Figure S7

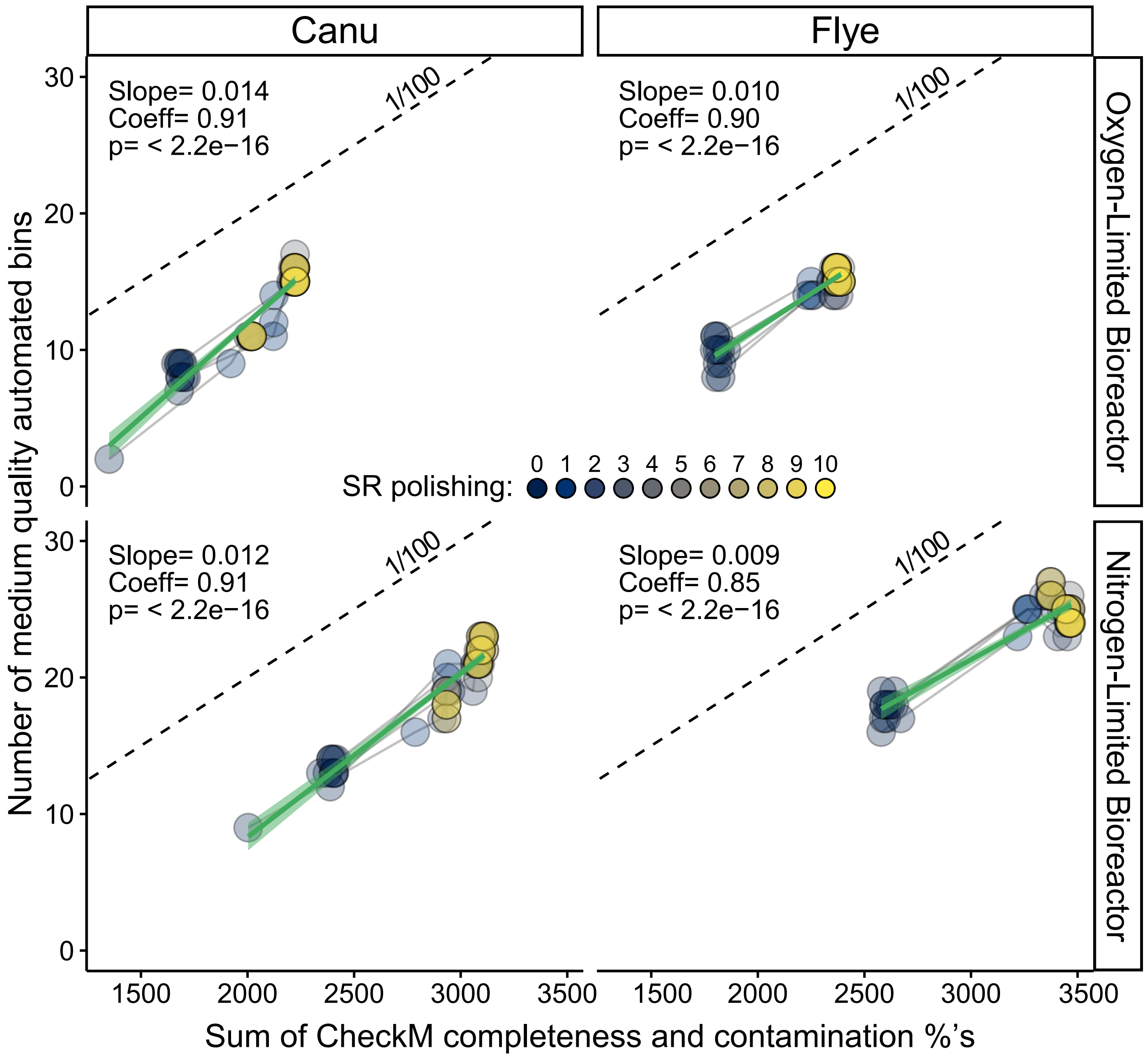


**Figure S7. Linear correlation between CheckM scores for an entire assembly and the medium quality or better (MQ) automated bin yield for each bioreactor and Long Read (LR) assembly throughout the iterative LR correction and Short Read (SR) polishing processes.**

The two bioreactors are separated over vertical panels, the two LR assemblers over the horizontal panels. Each point is colored by the SR polishing iteration, with grey lines connecting the points with the same number of preceding LR correction iterations, all of which are partially transparent. The green solid lines and shaded regions are the linear regressions for the displayed data and its 95% confidence interval. Broken gray line shows a slope of 1/100, representing the relationship that a redundancy score of 100 is theoretically equivalent to 1 MQ automated bin. Correlation coefficients (adjusted $R^2$) and p-values are displayed in the upper-left corner of each panel.
