## Supplementary material for "Simple, reference-independent analyses help optimize hybrid assembly of microbial community metagenomes": Figure S8

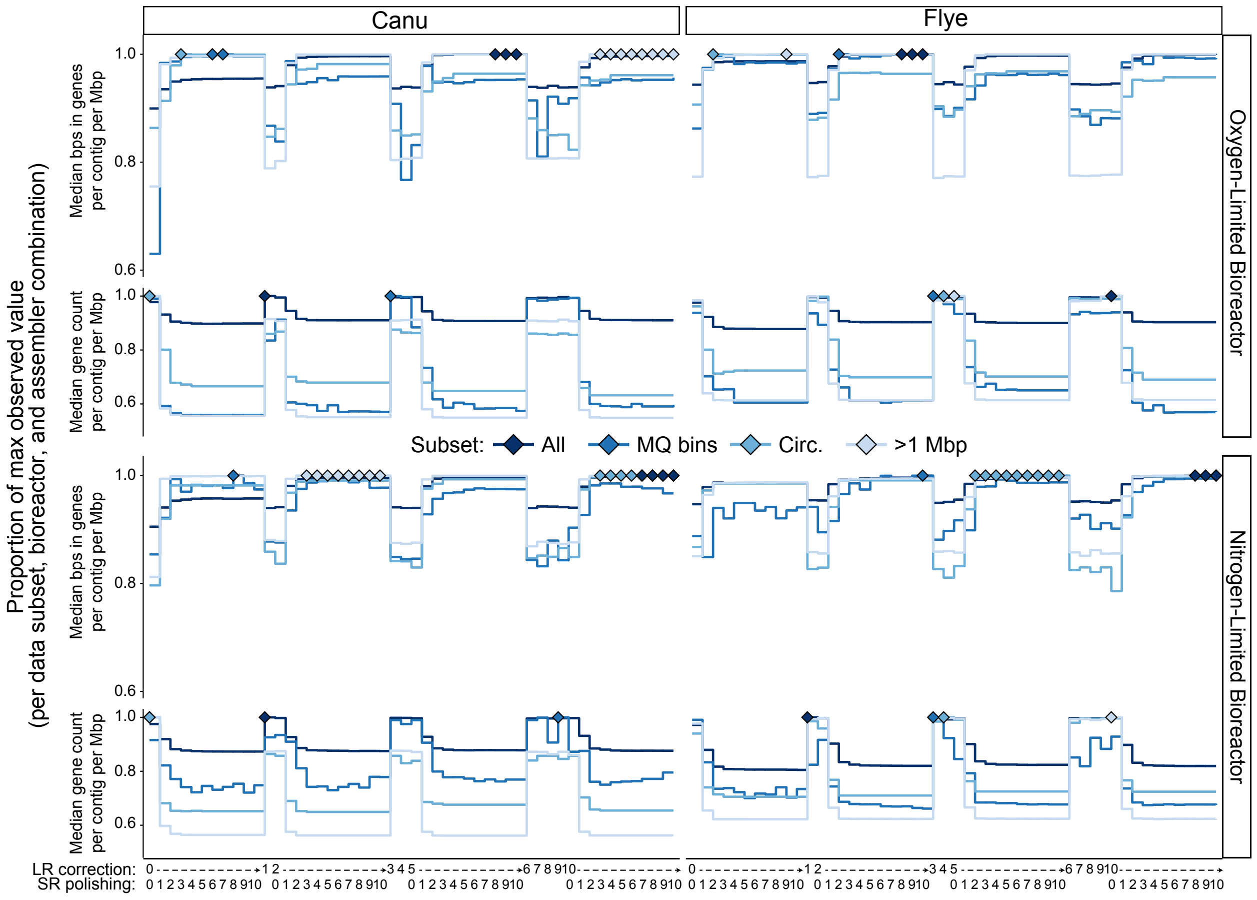


**Figure S8. Stair plot of predicted coding gene contents for each bioreactor and Long Read (LR) assembly throughout the iterative LR correction and Short Read (SR) polishing processes.**

The two bioreactors are separated over vertical panels, the two LR assemblers over the horizontal panels. The LR correction and SR polishing iterations are spread across the x-axis so that the ten SR polishing steps are immediately to the right of the preceding 0, 2, 5, or 10 LR correction step. Colored lines show the proportion of the maximum value for fractions of the assemblies throughout the LR correction and SR polishing iterations – the entire assembly (All), medium quality or better bins (MQ bins), circular contigs >10 Kbp (Circ.), and long contigs that may be complete but not circular bacterial genomes (>1 Mbp). Diamonds highlight the stages at which the maximum value occurred.
