## Supplementary material for "Simple, reference-independent analyses help optimize hybrid assembly of microbial community metagenomes": Figure S9

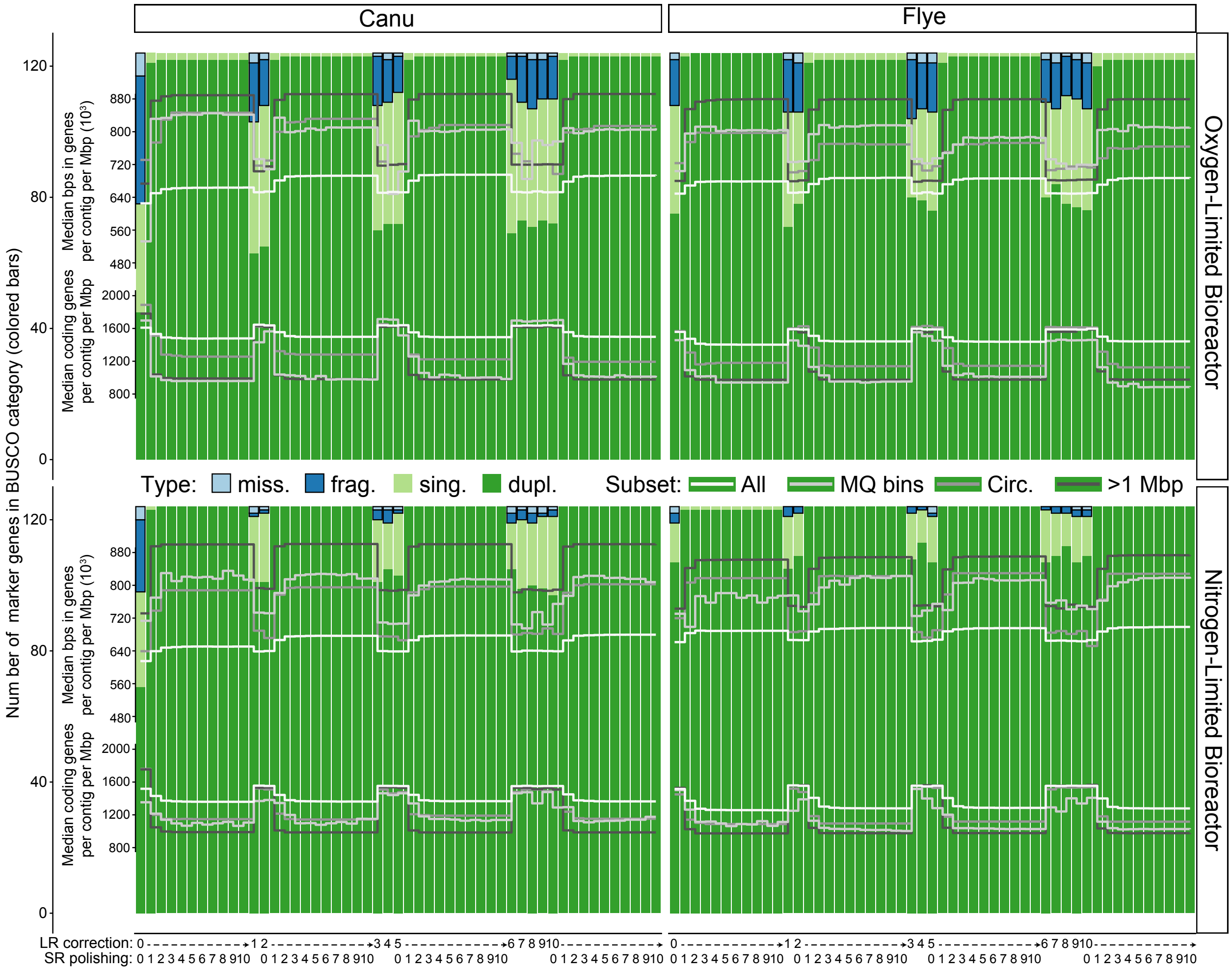


**Figure S9. Stacked bar plot of BUSCO marker gene recovery overlaid with stair plots of coding gene contents for each bioreactor and Long Read (LR) assembly throughout the iterative LR correction and Short Read (SR) polishing processes.** The two bioreactors are separated over vertical panels, the two LR assemblers over the horizontal panels. The LR correction and SR polishing iterations are spread across the x-axis so that the ten SR polishing steps are immediately to the right of the preceding 0, 2, 5, or 10 LR correction step. Colored bars show the number of bacterial marker genes in each BUSCO category – missing (miss.), fragmented (frag.), complete and single copy (sing.), complete and duplicated (dupl.) – for the entire assembly, and the gray lines indicate the median number of genes or bps in genes per contig per Mbp scaled to overlay the bars for several fractions of the assemblies throughout the LR correction and SR polishing iterations – the entire assembly (All), medium quality or better bins (MQ bins), circular contigs >10 Kbp (Circ.), and long contigs that may be complete but not circular bacterial genomes (>1 Mbp).
