## Supplementary material for "Simple, reference-independent analyses help optimize hybrid assembly of microbial community metagenomes": Figure S10

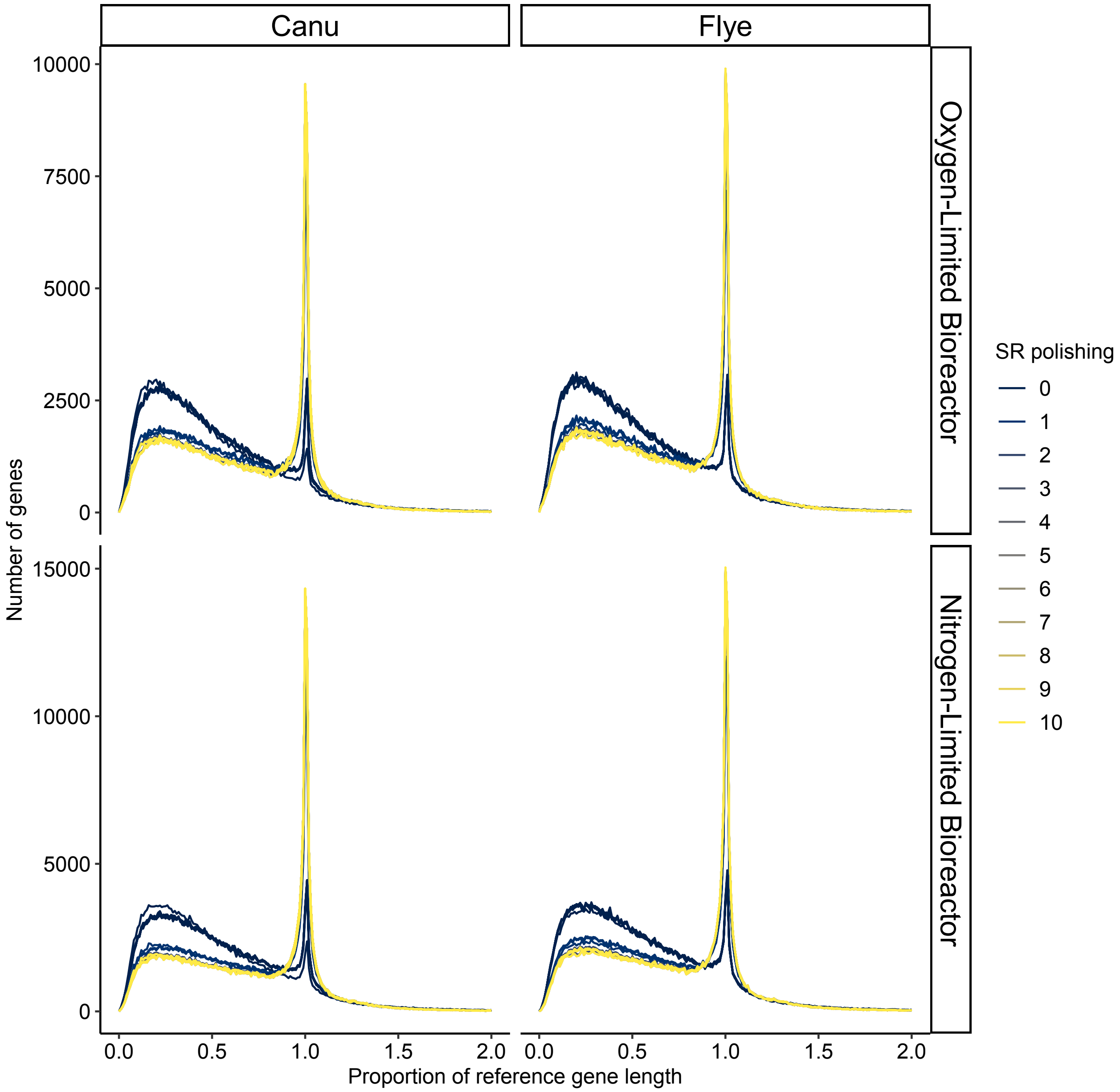


**Figure S10. Density curves of assembled gene lengths relative to their most similar reference determined using IDEEL for each bioreactor and Long Read (LR) assembly throughout the iterative LR correction and Short Read (SR) polishing processes.**

The two bioreactors are separated over vertical panels, the two LR assemblers over the horizontal panels. Colored lines indicate SR polishing iteration for the the gene sizes relative to their most similar reference sequence calculated by IDEEL for each assembly throughout the LR correction and SR polishing iterations. For clarity in the most relevant region, data are not shown for the proportion of reference gene lengths above two on the x-axis.
